# Fibroblast-derived thrombospondin-1 shapes macrophage polarization in advanced human co-culture models

**DOI:** 10.64898/2026.05.28.728363

**Authors:** Kristina Draganić, Sergey Isaev, Isabella Pototschnig, Daniel Valcanover, Janette Pfneissl, Mira Stadler, Marcos Hocevar, Vincent Lotz, Gabriel Wasinger, Céline Pisibon, Luisa Grader, Alina Sou Yan Ho, Małgorzata Sabina Małys, Piyal Saha, Renate Kain, Thomas Weichhart, Michael Bergmann, Walter Berger, Martina Schweiger, Igor Adameyko, Gerda Egger

**Affiliations:** Department of Pathology, Medical University of Vienna, Vienna, Austria; Department of Neuroimmunology, Center for Brain Research, Medical University of Vienna, Vienna, Austria; Institute of Molecular Biosciences, University of Graz, Graz, Austria; Center for Cancer Research, Medical University of Vienna, Vienna, Austria; Institute for Medical Genetics, Center for Pathobiochemistry and Genetics, Medical University of Vienna, Vienna, Austria; Department of General Surgery, Medical University of Vienna, Vienna, Austria; BioTechMed-Graz, Graz, Austria; BioHealth – Field of Excellence, University of Graz, Graz, Austria; Department of Physiology and Pharmacology, Karolinska Institutet, Stockholm, Sweden; Comprehensive Cancer Center, Medical University of Vienna, Vienna, Austria

**Keywords:** Tumor-associated macrophages, tumor microenvironment, lipid metabolism, organoids, fibroblasts, thrombospondin 1, CRC, co-culture models

## Abstract

**Background:** Tumor-associated macrophages (TAMs) are key drivers of the immunosuppressive tumor microenvironment (TME), supporting tumor progression through diverse functions. However, mechanistic studies of TAM polarization remain limited by the lack of physiologically relevant human model systems that capture stromal-immune interactions and macrophage heterogeneity.

**Methods:** We established advanced human co-culture systems that integrate healthy donor-derived macrophages with patient-derived organoids and tumoroids (PDOs and PDTs), as well as matched normal fibroblasts (NFs) and cancer-associated fibroblasts (CAFs). These multicellular models enabled the investigation of interactions among stromal, epithelial, and immune cells within tumor and adjacent normal tissue environments.

**Results:** The co-culture systems recapitulated distinct macrophage states associated with tumor and adjacent normal environments and identified fibroblasts as major regulators of macrophage phenotypes. CAFs promoted macrophage metabolic remodeling characterized by altered lipid handling and enrichment of TAM-like signatures. Mechanistically, we identified thrombospondin 1 (TSP1) as a CAF-secreted factor linked to metabolic priming. Recombinant TSP1 induced transient lipid accumulation followed by mitochondrial remodeling. In tumor co-culture conditions, CD36 inhibition reduced lipid accumulation in macrophages, supporting a role for TSP1-linked lipid crosstalk in stromal-immune interactions.

**Conclusion:** Our study establishes advanced patient-derived co-culture models as a platform to investigate human TAM biology and stromal-immune interactions in CRC. Using these systems, we identify a fibroblast-associated TSP1-lipid axis linked to macrophage metabolic remodeling and TAM-like polarization, highlighting stromal metabolic communication as a potential targetable feature of the CRC microenvironment.

**Graphical abstract:** 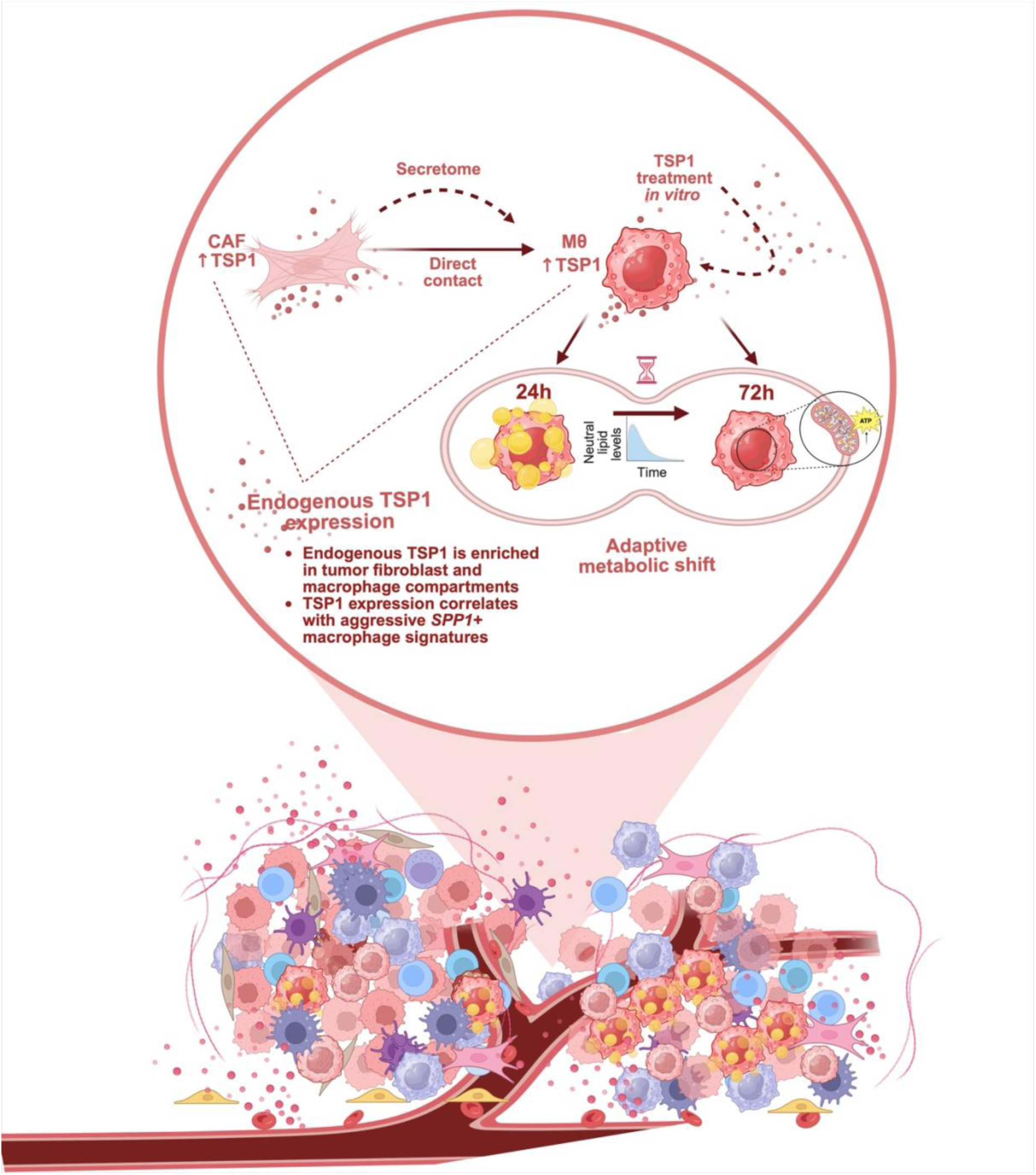

## Introduction

The tumor microenvironment has been recognized as one of the factors shaping cancer hallmarks. It is defined as a highly complex milieu in which cancer cells reside within the niches of different cell types that associate to form a specialized network, thereby shaping tumor physiology and therapeutic response (1,2). Therefore, the TME is a dynamic and adaptive ecosystem that fundamentally supports cancerogenesis, rather than merely acting as a passive bystander (3). Among the various cellular constituents of the TME, macrophages have emerged as a pivotal interface within this co-evolving environment and represent a major component (4,5). In the context of cancer, these cells are commonly referred to as tumor-associated macrophages and execute diverse tumor-promoting functions. TAMs actively support tumor growth by overcoming metabolic challenges and promoting inflammation, immunosuppression, angiogenesis, invasion, and metastasis (6–8). As a result, targeting protumorigenic TAMs has emerged as a promising strategy to disrupt their tumor-supportive role. Despite this potential, myeloid-targeted therapies that have entered clinical trials, such as anti-CSF1R agents, have demonstrated limited efficacy when used as monotherapies (9). The development of more effective treatments, as well as the identification of shortcomings in current approaches, relies on a deeper, more comprehensive understanding of TAM heterogeneity and function.

Traditionally, pre-clinical studies of macrophage biology have relied on two-dimensional (2D) cell lines, often examined within the M1/M2 polarization paradigm, as well as a variety of murine models (10,11). However, 2D cell lines possess substantial limitations: prolonged propagation, altered cell morphology, restricted polarization capabilities, and, critically, a lack of contact-dependent and environmental responsiveness to the TME, factors that are fundamental in shaping macrophage polarization. In addition, the binary M1/M2 classification has increasingly been recognized as an oversimplification of the true complexity of TAMs *in vivo*, with intermediate and diverse populations identified (12). Advances in single-cell technologies have further revealed an unprecedented molecular diversity among macrophages (13).

On the other hand, murine models have enabled the study of multifaceted cell–cell interactions in macrophage polarization in a more physiological setup. Nevertheless, significant challenges remain in extrapolating findings regarding ontogeny and metabolic profiles from murine to human macrophage populations, due to interspecies differences (14,15).

Therefore, the development of accurate and representative experimental models that capture not only the molecular heterogeneity of TAMs, as highlighted by single-cell analyses, but also their functional diversity is essential for advancing our understanding of macrophage biology and improving the efficacy of therapeutic strategies targeting TAMs.

Recently, the establishment of three-dimensional (3D) self-organizing stem cell-derived organoids has revolutionized patient-specific disease modeling. These organoids offer significant advantages over traditional two-dimensional (2D) models and have reduced reliance on animal experiments. Patient-derived organoids (PDOs) more accurately recapitulate the architecture and physiology of human tissues, providing a more relevant platform for studying cancer processes (10). Meanwhile, PDO platforms have evolved to include air-liquid interface (ALI) systems, microfluidic systems, and co-culture approaches (16). These advances enable the incorporation of additional TME components, further enhancing the physiological relevance of PDO models.

In this study, we developed a series of human co-culture models using colorectal cancer patient-derived tumoroids (PDTs), PDOs, fibroblasts, and healthy-donor PBMC-derived macrophages to create comprehensive platforms for studying human TAM polarization *in vitro*. Through these models, we focused on key signaling and metabolic cues that drive macrophage polarization toward pro-tumorigenic TAM phenotypes. Our findings revealed a lipid-mediated crosstalk between macrophages and fibroblasts and demonstrate that fibroblasts alone can prime macrophages into distinct subsets. Most notably, we identified the extracellular matrix protein TSP1, a CAF-derived secreted factor, as a mediator of macrophage priming toward pro-tumorigenic TAM subsets by altering their lipid metabolism profile. These insights not only illuminate the complex metabolic interplay within the TME but also highlight vulnerabilities that can be leveraged for therapeutic intervention.

## Results

### Establishment of advanced co-culture models to study macrophage biology

To capture human macrophage heterogeneity *in vitro*, study their polarization mechanisms, and enable efficient targeting of protumoral populations, we developed a series of co-culture models derived from primary human cells (Fig. 1A). Both epithelial and stromal components, including PDTs and CAFs, were isolated from colon cancer surgical specimens. From the same patient, we also generated NFs and PDOs from matched adjacent normal tissues. Macrophages were differentiated from monocytes isolated from peripheral blood mononuclear cells (PBMCs) of healthy donors and then co-cultured with fibroblasts (NF or CAF) in both direct and transwell co-culture models. Additionally, PDOs and PDTs from the same tissue sources were used in direct triple co-culture models, providing a comprehensive platform for interrogating macrophage-tumor-stroma interactions. Lastly, fibroblast-conditioned media were used to test the extent to which the fibroblast secretome alone could induce changes in macrophage phenotypes.

**Figure 1.**
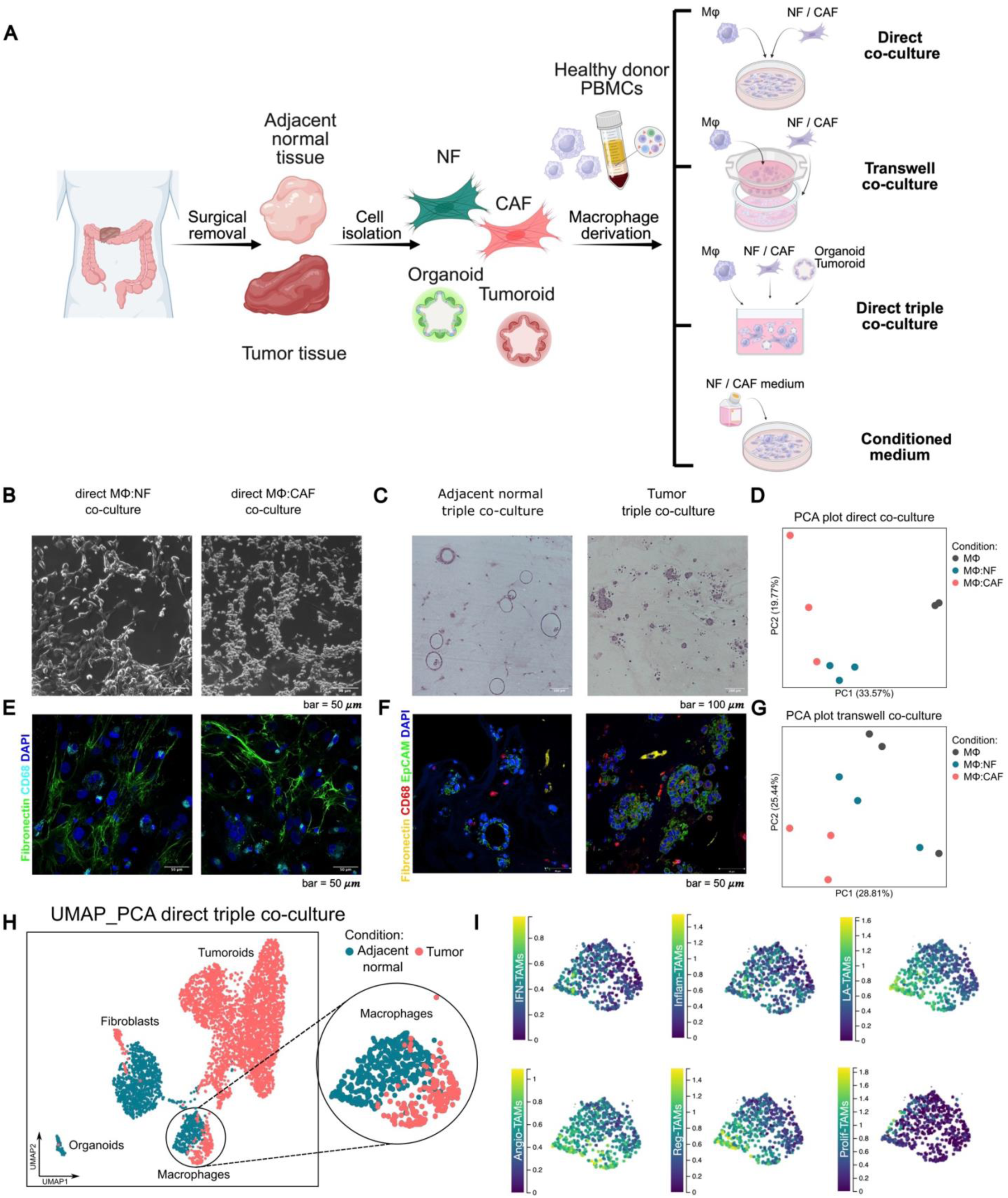
Establishment of co-culture models to capture human TAM heterogeneity. (A) Schematic representation of the co-culture configurations and cell origins used in this study. NF, normal fibroblast; CAF, cancer-associated fibroblast; MΦ, monocyte-derived macrophages. Created with BioRender.com (Draganic, K., 2026; https://BioRender.com/19btwmz). (B) Representative brightfield images of direct fibroblast-macrophage co-cultures, showing macrophages (MΦ) in normal fibroblast co-cultures (MΦ: NF, left) and cancer-associated fibroblast co-cultures (MΦ: CAF, right). (C) Representative brightfield microscopic images of sections of formalin-fixed paraffin-embedded (FFPE) triple co-cultures, stained with hematoxylin and eosin (H&E), containing adjacent normal patient-derived organoids (PDOs), NFs, and macrophages (left) and patient-derived tumoroids (PDTs) with CAFs and macrophages (right). (D, G) Principal component analysis (PCA) plots of bulk RNA-seq data of macrophages showing separation of macrophage populations between control (gray) and MΦ co-cultured with either NFs (green) or CAFs in (D) direct or (G) transwell co-culture models (n =3). (E, F) Representative confocal microscopic images of immunofluorescence staining with antibodies against Fibronectin 1 (FN1; fibroblast marker) and CD68 (macrophage marker) in (E) direct fibroblast-macrophage co-cultures or (F) additional EpCAM (epithelial cell marker) in triple co-cultures. (H) UMAP visualization of single-nuclei RNA-seq data from tumor versus adjacent normal co-culture models. Each point represents a single cell, colored by cell type annotation; macrophage populations are highlighted. (I) Enrichment of previously described distinct TAM signatures (13) in macrophage populations from the adjacent normal and tumor triple co-culture models, as in (C). TAM subset abbreviations: interferon-primed TAMs (IFN-TAMs), immune regulatory TAMs (Reg-TAMs), inflammatory cytokine-enriched TAMs (Inflam-TAMs), lipid-associated TAMs (LA-TAMs), proangiogenic TAMs (Angio-TAMs), and proliferating TAMs (Prolif-TAMs).

Microscopic images of the co-culture models illustrate extensive cell-cell interactions (Fig. 1 B, C, E, F, and Fig. S1A-D). Fibroblasts and macrophages formed interconnected network-like structures that were indistinguishable without cell-type-specific markers, such as CD68 for macrophages or fibronectin (FN1) for fibroblasts (Fig. 1E). Moreover, macrophages were attracted to PDOs, growing as hollow structures, and to tumoroids, showing dense growth patterns within the triple co-culture systems (Fig. 1C, F).

The extent of macrophage polarization across the various co-culture systems was assessed by transcriptomic analysis of macrophages isolated from fibroblast co-cultures (Fig. 1D, G). In co-culture with fibroblasts under both NF and CAF conditions, and in direct and transwell settings, significant changes in overall macrophage transcriptomic profiles were observed compared to macrophages in monoculture (Table S1). Additionally, all cell populations within the triple co-cultures were profiled at the single-nuclei level. The resulting UMAP clustering revealed separation of macrophage subsets between adjacent normal and tumor co-culture conditions (Fig. 3H). Importantly, heterogeneous macrophage populations were represented in the triple co-culture systems, including enrichment of various known pan-cancer TAMs (13) among macrophages in normal adjacent versus tumor triple co-cultures (Fig. 1I).

Finally, to demonstrate the potency of the fibroblast secretome, we analyzed the expression of classical pro- and anti-inflammatory markers in macrophages cultured in fibroblast-conditioned media (Fig. S1E). We observed an overall increase in CD11b expression. At the same time, CD86, a classical pro-inflammatory macrophage marker, was significantly decreased in macrophages cultured in both NF- and CAF- conditioned media. In contrast, the anti-inflammatory markers CD206 and CD163 were significantly upregulated, with a markedly stronger induction in CAF-conditioned macrophages. Interestingly, MHC class II expression was also increased in conditioned media–treated cells, whereas we observed no significant changes in SIRP α/β.

### Defining macrophage phenotypes in co-culture environments

To verify the relevance of our *in vitro* models in the context of CRC, we assessed the enrichment of *SPP1* and *C1QC* macrophage signatures (17) which represent two dichotomous, functionally distinct CRC macrophage subsets (18). In both direct and transwell co-cultures, we observed modulation of both signatures: while *SPP1* enrichment was higher in macrophages co-cultured with CAFs, *C1QC* enrichment was higher in those co-cultured with NFs (Fig. 2A, B). Moreover, a more pronounced effect was observed in this separation among macrophage populations in the triple co-culture model (Fig. 2C, D), where macrophages in the tumor conditions (PDTs plus CAFs) were strongly enriched for *SPP1*, and macrophages within adjacent normal conditions (PDOs plus NFs) were strongly enriched for *C1QC* gene signatures.

**Figure 2.**
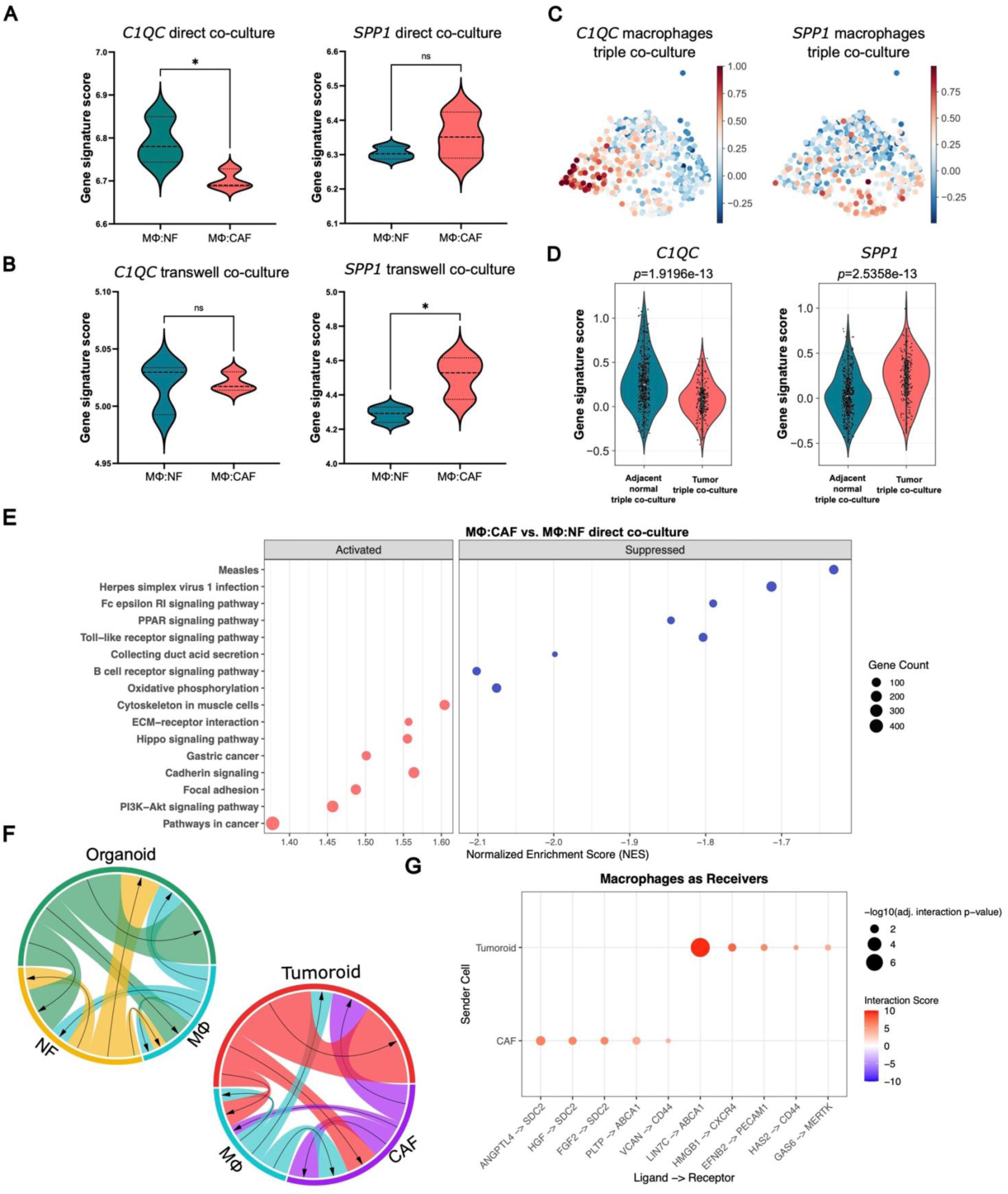
Molecular characterization and profiling of intercellular communication in macrophage subsets in co-culture models. (A, B) Panel showing enrichment of *C1QC* (left) and *SPP1* (right) gene signature scores (17) inferred from bulk RNAseq data of macrophages in (A) direct and (B) transwell co-culture systems. Data are plotted as boxplots representing gene signature scores across individual bulk macrophage samples under the indicated conditions (n = 3). Dashed lines indicate quartiles, and the central line represents the median. Statistical analysis was performed using a Student’s t-test (ns, not significant; \**p* < 0.05). (C, D) Distribution and quantification of enrichment for the same signatures in macrophage populations in adjacent normal and tumor triple co-cultures, inferred from single-nuclei RNA-seq data. Each dot represents an individual cell colored by enrichment of the indicated signature. Statistical analysis was performed using a Student’s t-test on per-cell score distributions. (E) Gene set enrichment analysis (GSEA) of the GO: KEGG database for direct MΦ: CAF vs. MΦ: NF co-culture transcriptomic comparisons, showing activated and suppressed pathways. Dot size indicates gene count, and color indicates directionality (NES, normalized enrichment score; activated: NES > 0; suppressed: NES < 0). (F) Chord diagrams summary of predicted ligand-receptor interactions from LIANA+ analysis (19) computed from single-nuclei RNA-seq data of tumor (top) and adjacent normal (bottom) triple co-cultures. Connection width reflects cumulative interaction score, and arrows indicate the predicted direction of signaling. Cell labels are harmonized for organoids (PDOs), tumoroids (PDTs), NFs, CAFs, and MΦ (macrophages) (G). Ligand-receptor interaction analysis of tumor co-cultures shows macrophages as communication receivers and predicted target genes involved in signaling with tumoroids or CAFs. Interactions were filtered for positive interaction scores and adjusted interaction *p* < 0.05. Color indicates interaction score, and size represents –log10 (adjusted interaction *p-*value).

Furthermore, pathway enrichment analysis of macrophages in CAF co-cultures revealed activation of cancer-related pathways, ECM-receptor interactions, cytokine-mediated, and TGF-β signaling (Table S2). In contrast, inflammatory pathways associated with viral and other infectious processes were suppressed (Fig. 2E, Fig. S2A).

In the triple co-culture setting, LIANA analysis (19) was conducted to infer cell–cell communication networks. The summary output indicated interactions among all cell types in both conditions (Fig. 2F, Table S3). Focusing on signals received by macrophages from CAFs in the tumor environment, *ANGPTL4*, *HGF,* and *FGF2* engaging *SDC2* and *VCAN* engaging *CD44* exhibited the strongest interactions, representing essential interactions that promote macrophage polarization toward immune-suppressive phenotypes. Interactions between PDTs and macrophages included *HMGB1-CXCR4, EFNB2-PEKAM,* and *GAS6-MERTK,* also driving M2-like macrophage polarization (Fig. 2G). Notably, the *ABCA1* receptor, an important cholesterol and phospholipid transporter, interacted with CAF-derived *PLTP* and tumoroid *LIN7C*, suggesting metabolic adaptations of macrophages in malignant conditions. In contrast, macrophages in the adjacent normal condition demonstrated higher scores for signaling axes initiated by collagens (*COL6A2*, *COL4A2*), *LAMC2*, *CXCL12,* or *CD59* engaging *CD44*, *CD151*, *CD4*, *STAB1*, or *TLR2* signaling axes, reflecting processes such as tissue homeostasis and baseline immune regulation (Fig. S2B)

Our co-culture data suggest that macrophage heterogeneity can be modeled *in vitro* using primary human cells and that fibroblasts are major constituents that prime macrophages toward distinct TAM subsets. To further investigate the contribution of fibroblasts, we profiled the surface marker expression of their secreted small extracellular vesicles (sEVs), which are well-established mediators of intercellular communication (20). As shown in the heatmap (Fig. S2C), sEVs derived from both CAFs and NFs were enriched for surface markers including CD29, CD142, CD40, CD146, CD105, CD44, CD49e, and MCSP. Notably, CD142, a factor involved in angiogenic and tumor-promoting processes (21), showed a tendency toward higher upregulation on CAF-derived sEVs (Fig. S2D). Moreover, CD29 and CD44 were both elevated in CAF sEVs. These receptors were shown to bind macrophage-derived SPP1, resulting in ECM remodeling and tumor cell migration. Moreover, direct crosstalk between macrophages and CAFs via these receptors shapes immune suppressive environments and resistance to immune checkpoint inhibitors (22,23). Together, these data highlight the potency of CAFs in shaping TAM phenotype through engagement with different surface receptors, either via direct cell-cell contact or via sEVs.

### Key stromal cues driving macrophage polarization in the tumor milieu

We next investigated the molecular pathways driving macrophage polarization in malignant conditions. Differential gene expression analysis comparing macrophages co-cultured with CAFs and NFs revealed consistent upregulation of *THBS1*, which encodes the extracellular matrix protein thrombospondin 1 (TSP1), in both the transwell and direct systems (Fig. 3A, B; Table S1). Furthermore, the *THBS1* gene signature (24) was strongly enriched in macrophages from triple co-cultures within the malignant environment (Fig. 3C). Upregulation of *THBS1* at the RNA level was confirmed in macrophages across all models (Fig. 3D). Notably, naive primary macrophages displayed lower baseline levels of *THBS1* expression. In contrast, exposure to CAFs, whether through direct or indirect contact or via the secretome, induced significant upregulation of this molecule, while NFs induced no significant induction. Further, upregulation of TSP1 was confirmed at the protein level in both co-culture with CAFs and in conditioned media (Fig. 3E, F, and Fig. S3A). Building on our previous findings from PDTs-fibroblast co-culture models (24), in which TSP1 influenced the fibroblast migratory phenotype, we next examined the expression profile of TSP1 in fibroblasts to determine whether they serve as a primary source of this molecule. Analysis of four patient-matched NF/CAF pairs revealed significantly elevated TSP1 levels at both the RNA and protein levels (Fig. 3G, H, and Fig. S3B).

**Figure 3.**
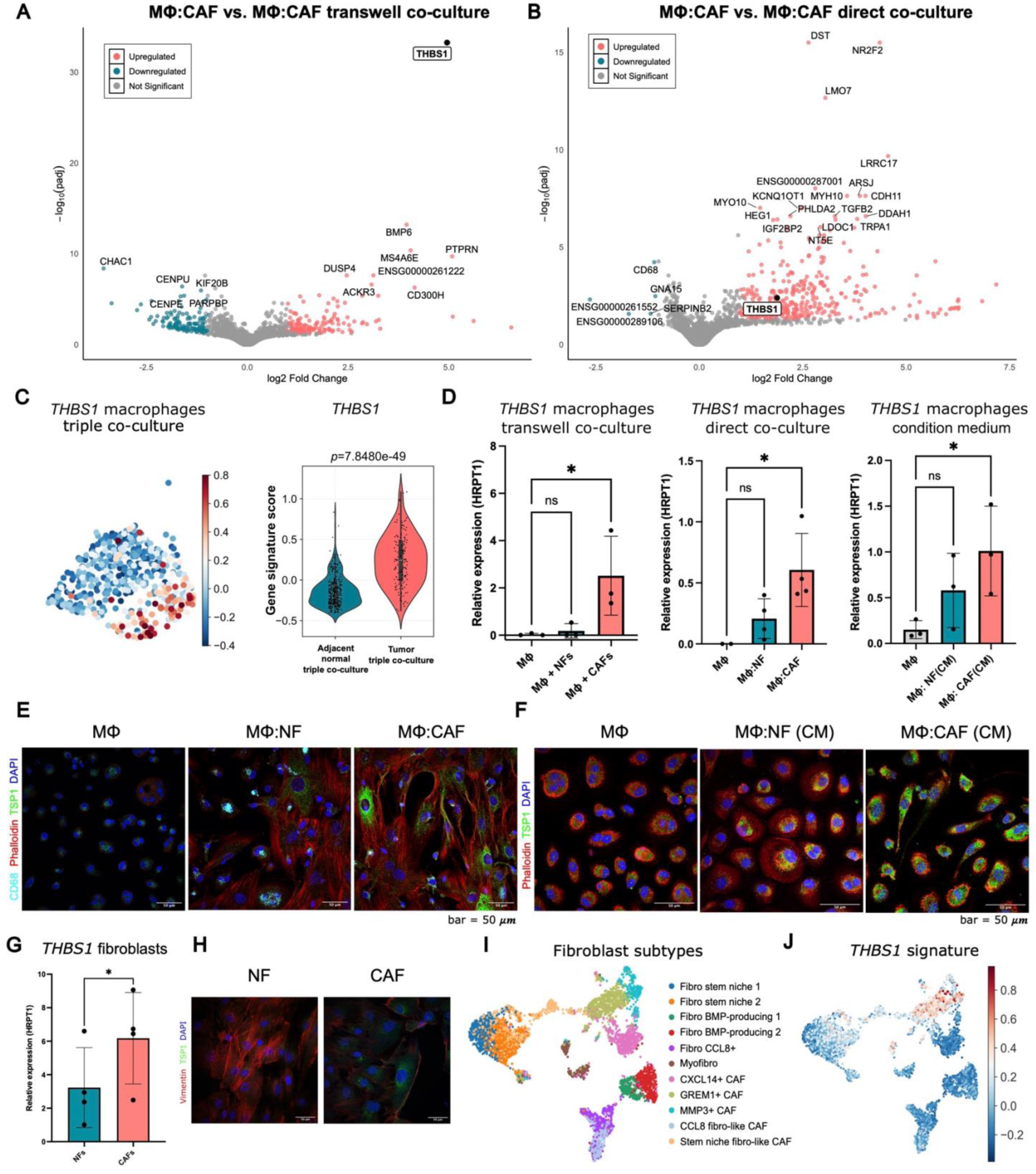
*THBS1* (TSP1) is upregulated in macrophages under co-culture in malignant conditions. (A, B) Volcano plots of differentially expressed genes between macrophages in CAF- versus NF-cocultures in (A) transwell and (B) direct co-culture models. Genes are classified as significantly upregulated (red), significantly downregulated (green), or not significant (gray) according to |log_2_FC| > 1 and adjusted *p* < 0.05; *THBS1* upregulation is highlighted. (C) Distribution of *THBS1* gene signature scores derived from single nuclei RNA-seq data of macrophages in triple cocultures, analyzed by Student’s t-test on per-cell score distributions. (D) mRNA expression of *THBS1* relative to *HRPT1* in macrophages cultured in transwell (n = 3), direct coculture (n = 4), or fibroblast conditioned media (n = 3). Statistical analysis by one-way ANOVA with Dunnett’s multiple comparisons test against monoculture controls (\**p* < 0.05; ns, not significant). (E, F) Representative confocal microscope images of immunofluorescence staining for phalloidin (red, F-actin), TSP1 (green), DAPI (blue), with CD68 (cyan, macrophage marker) additionally stained in co-culture conditions, showing (E) control macrophages, direct cocultures, MΦ: NF, and MΦ: CAF cocultures, and (F) control, NF, and CAF-conditioned media macrophages. (G) mRNA analysis of *THBS1* expression relative to *HRPT1* in tissue-matched NF versus CAF pairs (n = 4), analyzed by Student’s t-test (\**p* < 0.05). (H) Representative images showing TSP1 (green) expression in tissue-matched NF and CAF pairs (phalloidin, red; DAPI, blue). (I) UMAP representation of fibroblast subset clusters in a publicly available single-cell CRC dataset (25), with a corresponding plot showing the distribution of the *THBS1* gene signature (J).

To further characterize fibroblast populations expressing *THBS1* in patient samples, we reanalyzed a publicly available CRC single-cell RNA-seq dataset from Pelka et al. (25). Notably, we found that *GREM1+* fibroblasts, a population associated with *SPP1+* macrophage interaction (26), were enriched for the *THBS1* signature (Fig. 3I, J).

Finally, we examined the spatial distribution of the *THBS1* signature using spatial transcriptomics data from Faria de Oliveira et al. (27). Most macrophage and fibroblast populations, as identified by manual annotation, were found in the stromal regions, with fewer populations present within the tumor and WNT-high epithelial zone (Fig. S3C). Notably, the *THBS1* signature was primarily enriched in fibroblast and endothelial cell populations in the stroma. In addition, macrophages located near fibroblasts in these regions also showed a higher tendency for *THBS1* signature enrichment (Fig. S3D, E), emphasizing the spatial context of this signature’s expression.

### Relevance of TSP1 in the *in vivo* TME

We used a publicly available single-cell dataset from Pelka et al. (25) to analyze macrophage population clusters specifically. The distribution of *SPP1* and *C1QC* signatures in this patient COAD dataset matched previously reported patterns: *SPP1* was highly enriched in macrophages within tumor tissue, while *C1QC* was predominantly found in macrophages from adjacent normal tissue. We also observed a strong enrichment of the *THBS1* signature in tumor-associated macrophages (Fig. 4A, B). Notably, the *THBS1* signature showed a positive correlation with *SPP1* and a negative correlation with *C1QC*, suggesting that *THBS1*+ macrophages may represent a more aggressive population (Fig. 4C, D). Interestingly, this finding is further supported by evidence from RNA velocity analyses conducted by Qi et al., which inferred that the *SPP1*+ population may originate from *THBS1*+ macrophages (28).

**Figure 4.**
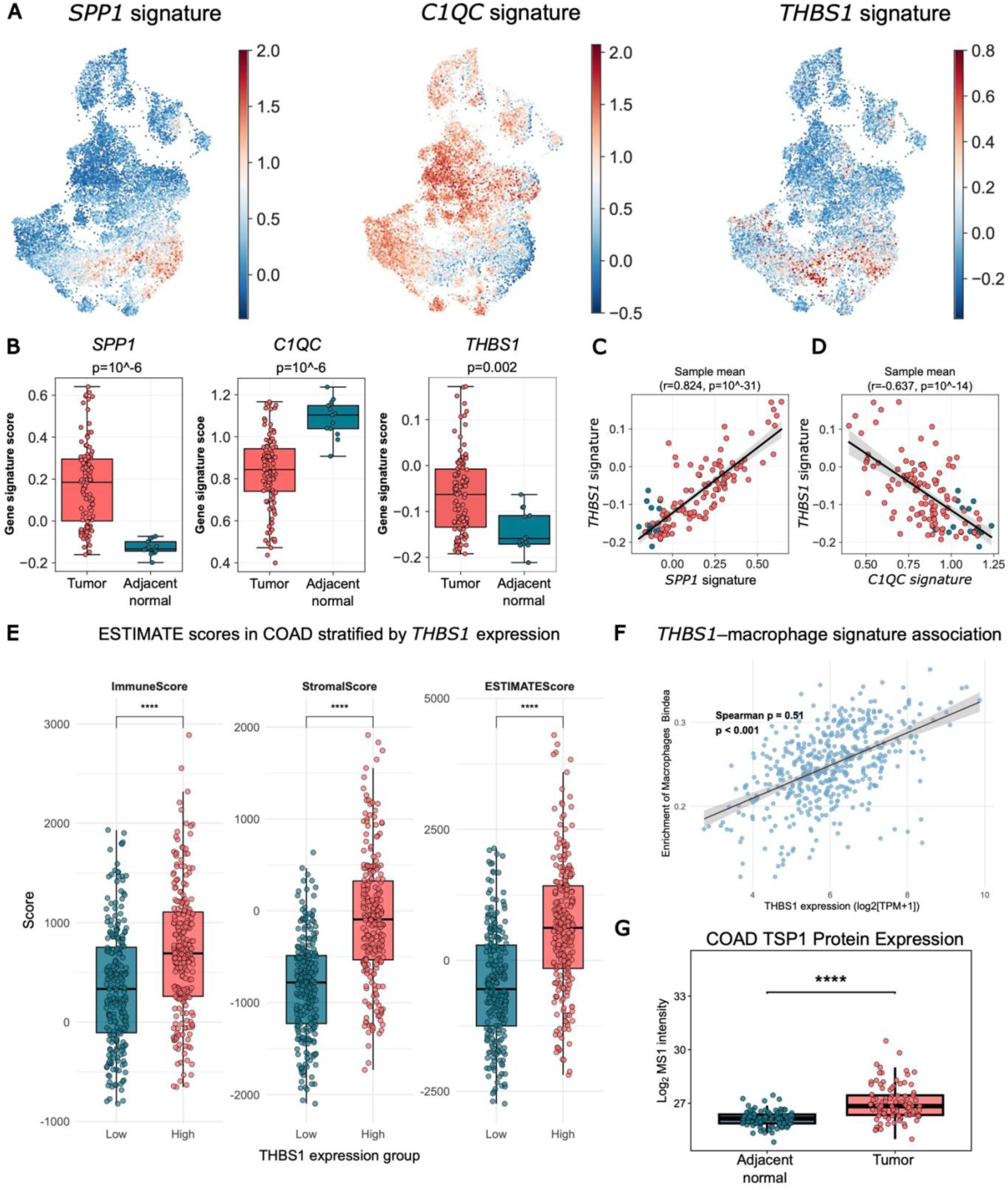
*THBS1* and *SPP1* signature scores are strongly correlated; *THBS1*-high tumors exhibit increased TME complexity. (A) Macrophage clusters in the Pelka et al. public dataset (25), colored by enrichment of *SPP1*, *C1QC*, and *THBS1* signature scores (17,47). (B) Quantification of signature score distributions, analyzed by Student’s t-test on per-cell scores. (C, D) Scatter plots showing correlations between (C) *THBS1* and *SPP1* and (D) *THBS1* and *C1QC* signature scores; correlation coefficients (r) and exact *p* values displayed. (E) ESTIMATE analysis (29) of the TCGA-COAD cohort (n = 481, Project ID: TCGA-COAD), stratified by *THBS1* median expression (*THBS1*-high, n = 240; *THBS1*-low, n = 241). Statistical comparison by Wilcoxon rank-sum test (\*\*\*\**p* < 0.0001). (F) Spearman correlation between *THBS1* expression (log₂[TPM+1]) and macrophage-related immune signature scores in TCGA-COAD (30) (n = 481) using Bindea et al. (54) macrophage gene set. (G) TSP1 protein levels in adjacent normal (n = 100) versus tumor (n = 97) COAD tissue from the CPTAC pan-cancer COAD dataset (31). Statistical comparison by Wilcoxon rank-sum test (*p* < 0.0001).

To extend our findings beyond the cellular level, we applied the ESTIMATE tool (29) to a TCGA COAD patient cohort (30) that had previously been stratified into *THBS1*-high and *THBS1*-low groups (Fig. 4E). Our goal was to assess TME complexity. In the *THBS1*-high patient group, we observed a significant increase in immune and stromal scores, resulting in a higher overall ESTIMATE score and indicating greater tumor complexity. Given this increase in immune cell infiltration, we further examined the relationship between *THBS1* expression and the immune context. As shown in Fig. 4F, *THBS1* expression was positively associated with the macrophage infiltration score in the patient cohort. These results further support the link between *THBS1* and a macrophage-enriched TME.

Considering that tumor tissue contains not only macrophages and fibroblasts but also other cell types, we assessed overall TSP1 protein levels in the COAD CPTAC cohort (31). In line with *THBS1*-associated gene programs observed in macrophages and fibroblasts, we found an overall increase in TSP1 protein expression in patient tumor tissue (Fig. 4G).

### Role of TSP1 in shaping macrophage metabolism and polarization within CRC TME

TSP1 has been described as an adipokine associated with obesity and non-alcoholic fatty liver disease (32,33). Mechanistically, TSP1 has been shown to support preadipocyte proliferation (34). Notably, lipid accumulation is a hallmark of TAM polarization and is associated with immunosuppressive and pro-tumorigenic functions (35). These observations prompted us to investigate whether exposure to recombinant TSP1 (rTSP1) could modulate the lipid profile of primary macrophages. To assess both early and late responses, we treated macrophages with rTSP1 at two concentrations (10 ng/mL and 1 µg/mL) and analyzed them at two different time points.

The early response (24h) to rTSP1 treatment resulted in significant accumulation of lipids in macrophages, as demonstrated by BODIPY™ 493/503 staining (Fig. 5A and B, and Fig. S4A and B). Representative images were selected to reflect the overall phenotype observed across male and female donors. Because of considerable inter-donor heterogeneity in baseline neutral lipid levels, we also quantified BODIPY™ 493/503 staining using FACS, which confirmed a significant increase in accumulation after treatment with TSP1 (Fig. S4C). However, prolonged exposure to rTSP1 (72h) reversed this effect, leading to a significant reduction in both spatially localized (Fig. 5C and D) and total lipid content (Fig. S4C).

**Figure 5.**
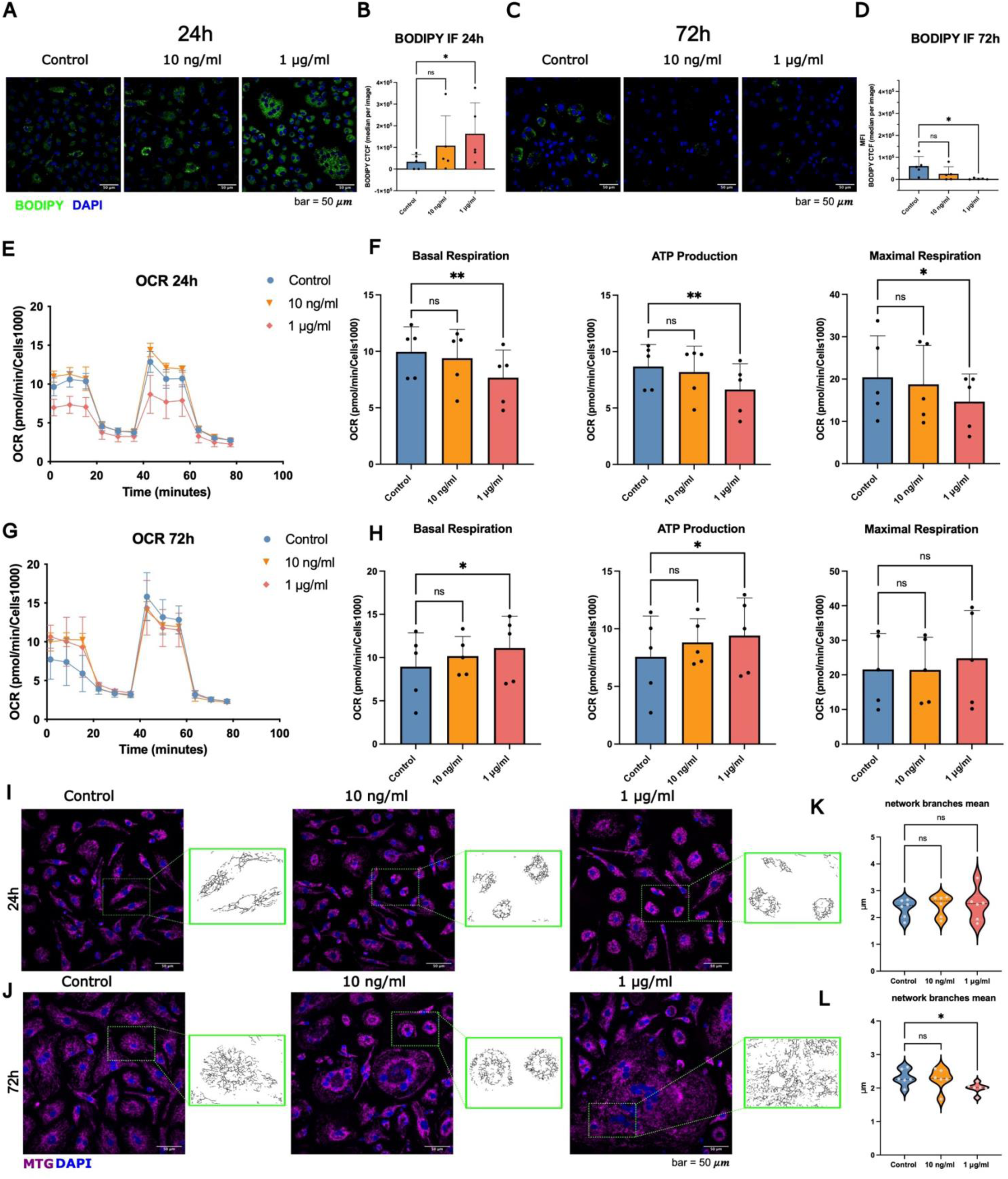
rTSP1 triggers early transient lipid accumulation and mitochondrial remodeling, followed by lipid clearance in macrophages. (A, C) Representative confocal images of BODIPY™ 493/503 staining in human male macrophages treated with control, 10 ng/ml, or 1 μg/ml recombinant TSP1 for 24 h (A) or 72 h (C). (B, D) Quantification of BODIPY staining, expressed as background corrected total cell fluorescence (CTCF), after (C) 24 hours or (D) 72 hours. (E, G) Representative Seahorse® MitoStress profiles showing oxygen consumption rates (OCR) of control (blue), 10 ng/ml (orange), or 1 μg/ml (red) TSP1 -treated macrophages after 24 h (E) or 72 h (G). (F, H) Quantification of indicated mitochondrial parameters from MitoStress tests at 24 h (F) and 72 h (H). Data shown in A-H panels are presented as mean ± SD from 5 donors. Statistical significance was determined using one-way ANOVA followed by Dunnett’s multiple comparisons test versus the control group (ns, not significant; \**p* < 0.05; \*\**p* < 0.01). (I, J) Representative MitoTracker Green (MTG) staining of the mitochondrial network in the indicated conditions at 24 h (I) and 72 h (J) in the male donor, with corresponding Fiji/ImageJ skeletonized images derived from the corresponding fluorescence images. (K, L) Quantification of mitochondrial network branches (mean value) from MiNA analysis (35) at 24 h (K) and 72 h (L). All data are shown as mean ± SEM from 6 donors. Statistical analysis was performed using one-way ANOVA with Dunnett’s multiple comparisons test (ns, not significant; \**p* < 0.05).

Given the transient lipid accumulation, we used Seahorse® analysis to investigate mitochondrial metabolism (Fig. 5E). The MitoStress test revealed a metabolically suppressed, quiescent mitochondrial state, with lower oxygen consumption rates (OCR) at 24 hours after higher dose rTSP1 treatment (Fig. 5F). rTSP1 treatment (1μg/ml) reduced basal respiration, ATP production, and maximal mitochondrial capacity. Additional parameters, including proton leak and spare respiratory capacity, were not affected, while non-mitochondrial oxygen consumption was reduced at high rTSP1 concentrations (Fig. S4D).

Conversely, lipid clearance at 72 hours post rTSP1 treatment was accompanied by a shift in the mitochondrial OCR profile at the higher treatment dose (Fig. 5G). This shift reflected mitochondrial functional reactivation, as indicated by increased basal respiration and ATP production. The trend toward maximal capacity was reversed compared to the early time point, although not significantly. Additional MitoStress-derived parameters showed increased proton leak, suggesting enhanced mitochondrial respiration and ATP production, and indicating metabolic remodeling with a partial uncoupling phenotype (Fig. S4E).

Following these observations, we sought to determine whether mitochondrial structural remodeling accompanied these dynamic metabolic changes. To this end, we performed MitoTracker Green (MTG) staining over time (Fig. 5I and J, and Fig. S4F and H). As highlighted by MiNa analysis (35), short-term TSP1 exposure at the higher dose led to a modest reduction in overall mitochondrial mass (Fig. S4). Other parameters reflecting mitochondrial complexity remained largely unchanged (Fig. 5K, Fig. S4G). In contrast, the metabolic shift toward increased mitochondrial activity during prolonged TSP1 treatment was associated with substantial structural remodeling, as evidenced by decreased branch length means and reduced network branching (Fig. 5L, Fig. S4I). Consistently, skeletonized images of orthogonal Z-stack projections (Fig. 5J, Fig. S4H) showed that the higher-dose treatment resulted in a marked decrease in mitochondrial interconnectivity.

Furthermore, mitochondrial structural and metabolic changes observed 72 hours after rTSP1 treatment were supported by specific changes in mitochondrial lipidomic profiles (Fig. 6A). While no significant overall changes in total lipid content between control and rTSP1 treated macrophages were evident (Fig. 6A), selected lipids specifically including cardiolipin (CL) and phosphatidylinositol (PI), were enriched following rTSP1 treatment (Fig. 6B, Fig. S5A, Table S4). Cardiolipins are essential for stabilizing the electron transport chain and maintaining mitochondrial structure. At the same time, phosphatidylinositol is a component of the outer mitochondrial membrane and implicated in endoplasmic reticulum (ER) crosstalk, mitochondrial dynamics, and lipid trafficking. Taken together, these results indicate a rewired mitochondrial state, potentially driven by early lipid-induced metabolic adaptations.

**Figure 6.**
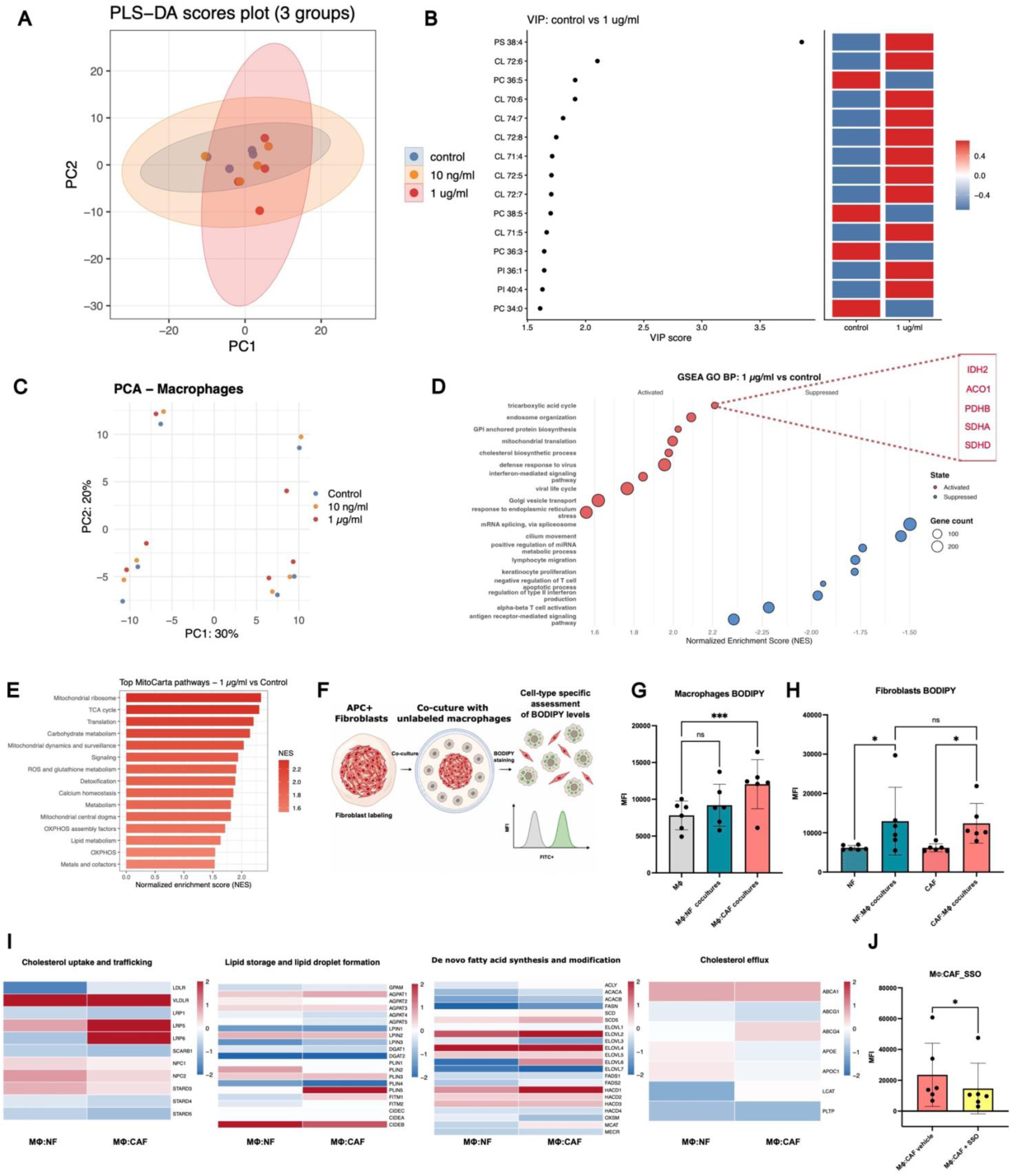
Recombinant TSP1 stimulation and co-culture both induce convergent lipid remodeling. (A) Partial least squares discriminant analysis (PLS-DA) score plot based on transformed and scaled normalized lipid abundances from the lipidomic profiling of human macrophages, visualizing the separation of control, 10 ng/ml, and 1 μg/ml rTSP1-treated groups 72 hours post-treatment (n=4). (B) Heatmap displaying variable importance in projection (VIP) scores for lipid species (VIP > 1.4) in 1 μg/ml rTSP1-treated versus control macrophages, as determined by pairwise PLS-DA models. PS, phosphatidylserine; CL, cardiolipin; PC, phosphatidylcholine; PI, phosphatidylinositol. (C) Principal component analysis (PCA) plot showing global transcriptomic changes in control, 10 ng/ml, and 1 μg/ml TSP1-treated macrophages after 72 hours. (D) GSEA of GO: BP pathways in macrophages treated with recombinant TSP1 (1 µg/ml) compared to control. Dot color indicates enrichment direction based on normalized enrichment score (NES): red for activated pathways (positive NES) and blue for suppressed pathways (negative NES), with dot size indicating gene count. Pathways involving TCA cycle genes are highlighted. (E) MitoCarta 3.0 pathway enrichment analysis of recombinant TSP1 (1 µg/ml)-treated macrophages versus control, with bar color indicating both magnitude and direction of enrichment. (F) Schematic illustration of BODIPY quantification in direct macrophage-fibroblast co-cultures, created in BioRender (Draganic, K., 2026; https://BioRender.com/ql19ozy). (G) BODIPY MFI measured by FACS in macrophage monoculture controls (MΦ), compared to MΦ: NF and MΦ: CAF co-cultured MΦ (n=6). Data are shown as mean ± SEM. Statistical analysis was performed using one-way ANOVA with Dunnett’s multiple comparisons test (ns, not significant; \*\*\**p* < 0.001). (H) BODIPY levels in fibroblast monocultures versus fibroblasts in co-culture (n=6). Statistical analysis was performed using one-way repeated-measures ANOVA followed by Šídák’s multiple comparisons test (ns, not significant; \**p* < 0.05). (I) Heatmap panel showing mean log2-transformed expression values for lipid metabolism-related genes in macrophages from direct MΦ: NF and MΦ: CAF co-cultures relative to those in monoculture. The color scale reflects the log2 fold change relative to MΦ. (J) BODIPY levels in macrophage-CAF co-cultured DMSO vehicle versus SSO-treated (50 μM). Statistical analysis was performed using a Student’s t-test (\**p* < 0.05).

Additionally, we profiled the bulk transcriptome of macrophages treated with low and high doses of rTSP1 under prolonged exposure using six biological replicates (Fig. 6C, Table S1). Although global transcriptomic changes were modest and showed pronounced inter-donor variability (Fig. S5B, C, D), GO: BP pathway enrichment analysis revealed upregulation of the TCA cycle, mitochondrial respirasome assembly, and aerobic respiration in both treatment groups (Fig. 6D, Fig. S5E, Table S2). These processes support mitochondrial structural and metabolic remodeling in response to initial lipid accumulation. Enrichment of mitochondria-related pathways, including TCA, OXPhos, and mitochondrial ribosome, was further confirmed using MitoCarta pathway analysis (Fig. 6E, Fig. S5F, Table S2).

Consistent with the modest overall transcriptomic changes, we observed a stable immunophenotype for classical pro- and anti-inflammatory macrophage markers following rTSP1 treatment (Fig. S5G). However, CD11b, an integrin commonly used as a general macrophage marker and known to negatively regulate anti-tumor responses (36), was significantly downregulated after rTSP1 exposure. Conversely, SIRPα/β, an inhibitor of phagocytosis (37), was upregulated. In line with this observation, we assessed macrophage phagocytic capacity using pHrodo™ BioParticles™ following rTSP1 treatment and found significant downregulation in the 1 μg/ml treatment group, compared with the control (Fig. S5H).

Altogether, our results show that exposure to rTSP1 caused only modest changes in gene expression and macrophages maintained a stable immunophenotype, but reduced macrophage phagocytic activity. Importantly, TSP1 triggered metabolic reprogramming, marked by early lipid accumulation, followed by mitochondrial structural and metabolic changes that ultimately promoted lipid clearance at later time points.

### Lipid-based macrophage–fibroblast crosstalk

To validate our findings on TSP1-mediated metabolic reprogramming in a more physiological context, in which CAFs supply TSP1, we studied lipid dynamics in a macrophage-fibroblast co-culture model. Before co-culturing, fibroblasts were pre-labeled with CellTracker Red to allow clear identification of each cell type (Fig. 6F). BODIPY staining revealed that lipid accumulation occurred specifically in macrophages co-cultured with CAFs (Fig. 6G). When CAFs were present, macrophages showed a significant increase in lipid content compared to monoculture controls. We also observed higher lipid content in co-cultured fibroblasts. However, this effect was not unique to CAFs, supporting the idea of reciprocal crosstalk in lipid metabolism between these cell types (Fig. 6H).

To further characterize these changes specifically in macrophages, we analyzed the expression of canonical lipid-associated markers involved in lipid scavenging and *de novo* synthesis and modification in our previously generated bulk transcriptomic dataset. Compared to controls, macrophages in co-culture showed upregulation of cholesterol-scavenging receptors (*LDLR*, *LRP5*, *LRP6*) and genes related to lipid modification and elongation (*SCD5*, *ELOVL2*, *ELOVL5*, *ELOVL6* (Fig. 6I). Additionally, genes involved in lipid droplet precursor formation and packaging (*AGPAT1*, *FITM1*, and *PLIN2*) were elevated in macrophages from CAF co-cultures. Finally, consistent with the upregulation observed in LIANA+ tumor triple co-culture macrophages, we found increased *ABCB1* expression, a cholesterol efflux transporter. To block lipid upregulation in MΦ: CAF co-cultures in a TSP1-dependent manner, we inhibited the TSP1 receptor CD36 using sulfosuccinimidyl oleate (SSO) (Fig. 6J). This treatment effectively reduced lipid accumulation in macrophages co-cultured with CAFs and in the CAFs themselves (Fig. S5I). Notably, we did not see significant changes in neutral lipid levels in macrophage monocultures, NF co-cultured macrophages, or normal fibroblasts co-cultured with macrophages, emphasizing the specific importance of CD36-mediated lipid uptake in the tumor context (Fig. S5J-L). Our results suggest that TSP1 in co-cultures may be a key factor in maintaining a lipid-rich macrophage-CAF interaction state.

## Discussion

Recent advances in single-cell technologies have revealed the remarkable heterogeneity of tumor macrophage populations (13). However, efforts to define the functional significance of these diverse subsets have been limited by a lack of robust *in vitro* models that faithfully capture this complexity, support in-depth functional studies, and aid the development of therapies targeting protumorigenic TAMs. Here, we developed patient-derived co-culture models to study TAM polarization and the mechanisms of myeloid-driven immunosuppression. Incorporating macrophages, tissue-matched primary fibroblasts, and PDOs or PDTS, these models closely recreate the complexity of the CRC TME, allowing direct comparison of tumor and adjacent normal tissue. They overcome the limitations of traditional monocultures for exploring stromal-immune-tumor interplay and enable the investigation of tumor-specific processes.

Our co-culture models demonstrate that CAFs play a pivotal role in shaping macrophage niches and priming naive macrophages into distinct TAM subsets via direct contact or through their secretome. This supports recent reports identifying spatial *FAP+* fibroblast subsets and *SPP1+* macrophages at the tumor sites, underscoring both the structural and functional significance of this stromal-immune crosstalk (28).

We identified a key metabolic hallmark of macrophages interacting with CAFs: substantial accumulation of neutral lipids. This phenotype may be partly explained by enhanced scavenging activity, as macrophages in CAF co-cultures upregulate genes involved in cholesterol uptake and lipid processing, further supporting previously described mechanisms of lipid-associated macrophage generation (38,39). However, alongside neutral lipid accumulation, we observed an increase in cholesterol efflux transporters, suggesting a compensatory mechanism that is simultaneously active in response to increased intracellular lipid storage. In contrast, fibroblasts exhibited a broad increase in lipid levels regardless of activation status, highlighting distinct metabolic dynamics between stromal and immune compartments. This observation is consistent with a study reporting significant enrichment of diverse lipid classes in conditioned media, but not in intracellular compartments, in CAFs compared with NFs (39,40). Nevertheless, we speculate that the CAF secretome could also contribute to the lipid pool that promotes macrophage scavenging.

While lipid metabolism in macrophages is known to promote immunosuppression and tumor progression across cancers (38,41), our study uncovers a distinct axis of macrophage–fibroblast lipid crosstalk. Previous research, such as that by Kloosterman et al., has focused on macrophage-mediated myelin degradation in glioblastoma and on macrophage-derived cholesterol in prostate cancer therapy resistance (42,43). Additionally, epigenetically dysregulated PDAC cancer cells influence the lipid phenotype of CAFs (44). By highlighting the interplay and spatial proximity of macrophages and fibroblasts, our findings shift attention from tumor cell-centric mechanisms to the critical role of stromal–immune interactions. This novel perspective opens new avenues for therapeutic intervention targeting the tumor microenvironment.

We furthermore pinpointed TSP1 as a central signaling molecule in the crosstalk between macrophages and CAFs. TSP1, an ECM protein, has been classically described as an angiogenesis inhibitor, though its role in tumorigenesis is highly context-dependent (45). More recently, its role in immunity and TAM biology has been reevaluated, with accumulating evidence indicating that *THBS1*-expressing macrophages serve as precursors of the *SPP1+* macrophage subset (28). Consistent with these insights, our data revealed a strong correlation between these gene signatures in patient-derived macrophages, providing further evidence for this connection. *THBS1+* monocytes and myeloid populations have also been connected to metastatic behavior and immunosuppression in both colorectal and hepatocellular carcinoma (46,47). While previous studies primarily described an indirect role for TSP1, such as promoting cytotoxic T-cell exhaustion and impairing vascularization in mouse models of colorectal cancer (46), our investigation focused on the direct effects of TSP1 on human macrophage function and metabolism, specifically in the context of lipid metabolism, since previous work has established a role for TSP1 in forming lipid-rich lesions in a leptin-mediated atherosclerosis model (48). To mimic conditions where CAFs secrete TSP1, we used recombinant TSP1 at two concentrations: one approximating physiological blood levels in healthy donors (4.5–45 ng/ml) and a higher dose representing pathologically elevated serum concentrations observed in colorectal cancer patients (18–20 μg/ml) (46,49). Our recombinant protein experiments showed that acute TSP1 treatment led to a marked but transient accumulation of neutral lipids in macrophages. Notably, after 72 hours of TSP1 exposure, we observed enhanced lipid clearance accompanied by mitochondrial structural and metabolic changes. These included elevated basal respiration and increased ATP production, characteristic of a lipid-oxidative, tumor-promoting macrophage phenotype. Such metabolic effects were prominent at the higher TSP1 dose, while gene expression changes at the lower dose indicated a similar direction. Transcriptomic analysis revealed upregulation of mitochondrial pathways, including the TCA cycle, oxidative phosphorylation, and mitochondrial translation. Lipidomic analysis showed increased cardiolipin levels. Given the previously described TSP-CD47 and CD36-mediated signaling and their association with mitochondrial ROS generation and dynamics (50,51), the observed increase in cardiolipin levels, together with alterations in mitochondrial structure, likely reflects adaptive remodeling. These changes appear to support enhanced oxidative respiratory capacity, indicating a metabolic adaptation rather than overt mitochondrial dysfunction.

Interestingly, pathway analysis highlighted a reduction in macrophage immune programs associated with T-cell activation following TSP1 treatment, consistent with prior findings indicating that TSP1 can directly inhibit T-cell activation via CD3/CD28 pathways (46). Beyond these metabolic shifts, we observed functional changes in macrophages, specifically reduced phagocytic capacity, which is essential for anti-tumor immunity. Importantly, these functional alterations occurred without substantial shifts in the expression of conventional pro- and anti-inflammatory markers, further reinforcing the notion that TAM polarization extends beyond the traditional M1/M2 classification scheme.

Finally, we managed to reverse lipid-mediated crosstalk under physiologically relevant conditions in a TSP1-dependent context. By employing SSO, a direct competitor of TSP1 for CD36, we specifically reduced neutral lipid levels in both macrophages and CAFs within tumor-specific co-cultures. These findings underscore the critical importance of TSP1-CD36-mediated lipid uptake in the macrophage co-culture model.

In summary, our study offers a new framework for investigating TAM biology in human-relevant systems and uncovers therapeutic opportunities targeting stromal lipid metabolism. While dietary fat and obesity are established CRC risk factors (52), and lipid-pathway-targeting agents have already been recognized for their potential to reduce CRC mortality (53), our findings extend this concept by showing that targeting the metabolic support provided by the stroma may further enhance responses to combination cancer therapies.

## Conclusion

This study highlights the central role of lipid-based intercellular communication and stromal cues in macrophage polarization within the CRC TME, particularly through TSP1-mediated effects. Competition for the CD36 receptor modulates lipid levels and mitochondrial activity in macrophages, potentially generating metabolic vulnerabilities and presenting a strategy for macrophage reprogramming.

## Limitations

While our human co-culture system offers advantages over mouse models and cell lines, it still relies on non-autologous macrophages from healthy donors, which introduces significant donor-to-donor variability. We anticipate that if we used a fully autologous system, polarization effects might be even stronger. Our model also lacks key components of the TME, such as endothelial cells and T cells, which could further affect cell-cell communication. For instance, in the spatial analysis of CRC patient data, we observed an increase in the TSP1 signature in endothelial cells. We also did not explore how macrophages affect NFs/CAFs and PDOs/PDTs, since our focus was on macrophage biology. Another limitation was the reliance on pharmacological treatment (e.g., rTSP1) of terminally differentiated primary macrophages, which are difficult to modify genetically.

We also recognize that the tumor cytokine environment is highly complex, with many factors that may interact with TSP1 in both synergistic and antagonistic ways. Another challenge is that current methods for targeting TSP1 are indirect, as they inhibit its receptors on cells rather than TSP1 itself. Future research should prioritize testing these interactions and developing more selective inhibitors to assess their impact better.

## Material and methods

### Generation of human peripheral blood mononuclear cell (PBMC)-derived macrophages and rTSP1/SSO stimulation

Human macrophages were generated from whole-blood samples of healthy donors. Monocytes were isolated from PBMCs by density gradient centrifugation with LymphoPrep (StemCell Technologies). Cells were plated in ultra-low attachment dishes (Corning) for 3 hours in FBS-free DMEM (Gibco) at 37°C with 5% CO2. Adherent cells were cultured in RPMI 1640 medium with 10% FBS (Gibco), 1% Penicillin-Streptomycin (Gibco), and 20 ng/mL M-CSF (PeproTech) for differentiation, with one medium switch on day 3. Fully differentiated macrophages were used for experiments on day 6 and detached with Accutase (Gibco). For the recombinant protein study, human rTSP1 (R&D Systems, cat. No. 3074-TH) was used at the concentrations indicated in the graph. Sulfosuccinimidyl oleate (SSO; MedChemExpress, cat. no. HY-112847A) was used at a final concentration of 50 μM for 72 hours, with one media replenishment.

### Cultivation of patient-derived organoids (PDOs), patient-derived tumoroids (PDTs), and cancer-associated fibroblasts/normal fibroblasts (CAFs/NFs)

PDOs and PDTs were established from tissue-matched adjacent normal or tumor samples as previously described (24,55). PDTs were maintained in ENAS medium (Table S5), and PDOs in WENRAS medium, which is ENAS supplemented with WNT3a and R-spondin as previously described (56). Cells were passaged every 7–10 days with TrypLE (Gibco). Primary fibroblasts were isolated from the same specimen (NFs from adjacent normal and CAFs from tumor tissues), propagated in Endothelial Cell Growth Medium (PromoCell), and passaged every 3 days using Trypsin (Gibco). All lines used in this study were in passages P3–P12. The mutational profile of the PDOs/PDTs used in this study is listed in Table S5.

### Double and triple co-culture models

Double co-cultures were prepared using two methods: direct co-culture and transwell co-culture. For direct co-cultures, fibroblasts, CellTracker-labeled (Thermo Fisher Scientific, cat. No. C34565, according to the manufacturer’s protocol) or unlabeled, were mixed with macrophages at a 1:2 ratio. In the transwell model, fibroblasts were embedded in a 3D matrix composed of 30% Matrigel (Corning) and 70% Collagen I (Corning) at the bottom of the well. At the same time, macrophages were seeded in a transwell insert (Corning) placed above the matrix. Additionally, macrophages were mixed with CAFs or NFs, and with PDTs or PDOs, in direct co-cultures within the same matrix. Before co-culture, PDTs and PDOs were seeded as single cells and allowed to form structures over 2–5 days in their respective maintenance media. They were then differentiated in Human IntestiCult Org Diff Medium (StemCell) for 5 days, with two medium changes. For harvesting, direct co-cultures were detached using Accutase (Gibco) and mechanical scraping, whereas matrix-embedded cultures were digested using TrypLE (Gibco) and Collagenase B (Roche). For separation from fibroblasts in the direct co-culture model, the co-culture was harvested as a pellet and incubated with human CD11b MicroBeads (Miltenyi, cat. no. 130-097-142) according to the manufacturer’s protocol. After specific removal of fibroblasts, the isolated cells were used for downstream applications. Conditioned media were collected after culturing fibroblasts in Human Plasma-Like Medium (HPLM) for 3 days. The supernatant was filtered through a 0.2 µm filter and diluted 1:1 with fresh HPLM. All co-cultures were maintained in HPLM (Gibco) supplemented with 10% FBS (Gibco) and 1% Penicillin-Streptomycin (Gibco).

### Immunofluorescence and immunohistochemistry staining

For microscopic imaging, direct co-cultures were seeded in µ-Slide 8-well chambers (ibidi) and maintained under standard culture conditions. Cells were washed with 1× PBS and fixed with 4.5% paraformaldehyde (PFA) for 10 min at room temperature. Following fixation, cells were washed with washing buffer (0.1% Triton X-100 and 0.05% Tween 20 in 1× PBS), permeabilized using permeabilization buffer (0.5% Triton X-100 in 1× PBS), and subsequently blocked with blocking buffer (10% goat serum and 0.1% Triton X-100 in 1× PBS). Primary antibodies were diluted in blocking buffer and incubated overnight at 4°C, then washed with washing buffer and incubated with appropriate secondary antibodies or a staining solution listed in Table S5. Nuclei were counterstained with DAPI. Triple co-cultures were processed for paraffin embedding and imaged as tissue sections. Briefly, matrix-containing cultures were fixed with 2% PFA for 30 min, embedded in paraffin, and sectioned into 3 µm-thick slices, mounted on microscope slides. For immunofluorescence staining, paraffin sections were deparaffinized, subjected to microwave-based antigen retrieval, blocked with 10% goat serum, and incubated overnight at 4°C with primary antibodies. Sections were subsequently washed with TBS, then incubated with the appropriate secondary antibodies and DAPI counterstaining (Table S5). All images were acquired using an Axio Imager M2 microscope (Zeiss) with ZEN 2 Blue Edition software (Zeiss). Triple co-culture images were generated by overlaying images acquired from two consecutive sections. For hematoxylin staining, slides were deparaffinized at 56 °C for 30 min and stained with hematoxylin (Sigma-Aldrich) for 5 minutes. Slides were then mounted using mounting medium (Aquatex), sealed with coverslips, and allowed to dry.

### Image-based fluorescence quantification (CTCF)

Fluorescence intensity was measured using Fiji software (ImageJ, v1.54p). DAPI-stained nuclei were segmented to define c). For each ROI, area, mean intensity, and integrated density (IntDen) were recorded. A background ROI was manually selected for each image, and its mean intensity was used to correct the background. Corrected Total Cell Fluorescence (CTCF) was calculated for each ROI as IntDen − (Area × background mean). CTCF values were determined for each cell in every image and summarized as the median CTCF per image. Each donor was imaged in triplicate. Image-level median values were plotted and used for downstream statistical analysis. Statistical analyses were performed according to the experimental design using either Student’s *t*-tests or one-way ANOVA with Dunnett’s multiple comparisons correction.

### Single-Nuclei Library Preparation

Snap-frozen samples were processed to generate single-nuclei ATAC and gene expression libraries using customized 10X Genomics protocols. Nuclei isolation was performed using the Chromium Nuclei Isolation Kit with RNase Inhibitor (10xGenomics, PN-1000494) according to the user guide CG000505, Chromium Nuclei Isolation Kit, UG, RevA, with slight modifications. In brief, 200 µl of lysis buffer was added to the sample, which was then dissociated with the vendor-provided pestles. The sample was further homogenized by pipette-mixing and incubated on ice for 10 min. Afterward, the sample was washed twice with 1ml of Wash Buffer, followed by centrifugation for 5min at 500 rcf at 4°C. For multiplexing, a customized approach (Schwarz et al., in revision) based on barcoding with cholesterol-modified oligos (CMOs) was applied to pool up to 4 samples per single-nuclei sequencing reaction. For this, nuclei were subsequently incubated with unique CMO Anchor-Barcode and Co-Anchor, and thereafter washed twice with 1 ml of Wash Buffer by centrifugation at 500 rcf for 5 min at 4°C. The final nuclei pellet was taken up in 30 µl Diluted Nuclei Buffer (10X Genomics). Nuclei were counted using Acridine Orange/Propidium Iodide Stain (Logos, F23001) and an automated dual fluorescence cell counter (Logos, LUNA-FL™). Equal numbers of nuclei were pooled per multiplexed reaction, resulting in a total maximum of 16,000 nuclei per individual Chromium Next GEM Single Cell Multiome ATAC + Gene Expression (10X Genomics, PN-1000285) reaction. The pooled nuclei were transposed, loaded on a Chromium Next GEM Chip J (10X Genomics, PN-1000230), and run on the Chromium iX instrument as instructed by the manufacturer. ATAC and gene expression libraries were generated according to the user guide CG000338, Chromium Next GEM Multiome ATAC GEX User Guide RevG. The CMO hash library was generated by adding an additive primer to the cDNA amplification, purifying hash-DNA from the supernatant during cDNA cleanup, and ultimately amplifying CMO hash libraries with unique i5 and i7 index combinations. Library quality and size were assessed with the High Sensitivity DNA Analysis Kit (Agilent Technologies #5067-4626) and the Bioanalyzer 2100. For sequencing, gene expression libraries were pooled with 17% corresponding CMO hash libraries and sequenced on the Illumina NovaSeq6000 system using read 1: 28 cycles, i7: 10 cycles, i5: 10 cycles, and read 2: 90 cycles. The ATAC libraries were sequenced on the Illumina NovaSeq6000 system using read 1N: 50 cycles, i7: 8 cycles, i5: 24 cycles, and read 2N: 49 cycles.

### Analysis of scRNA-Seq

Raw sequencing reads were processed with Cell Ranger ARC v2.0.0 (10x Genomics). Samples had been pooled prior to sequencing using CellPlex multiplexing, in which each sample was labeled with a distinct CMO, captured alongside the transcriptome, and used to assign each cell barcode to its sample of origin. Four CMOs were used in this experiment, with the following barcode sequences: ACCCACCAGTAAGAC (adjacent normal tissue), GGTCGAGAGCATTCA (tumor tissue), CTTGCCGCATGTCAT (adjacent normal triple co-culture), and AAAGCATTCTTCACG (tumor triple co-culture). Demultiplexing assigned each cell to one of these four samples based on the dominant CMO signal. Of the four samples, only the adjacent normal and tumor triple co-cultures populations passed quality control and were retained for downstream analysis; the healthy and tumor tissue samples were excluded due to insufficient cell recovery and poor QC metrics. Following demultiplexing, cells called by Cell Ranger’s default filtering were further filtered to remove low-quality barcodes and likely doublets. Further data analysis was performed using the Scanpy (v1.11) Python package (57). Cells were required to have less than 20% of reads mapping to mitochondrial genes (a marker of cell stress or lysis), at least 1,000 detected genes (to exclude empty or poorly captured droplets), and a Scrublet doublet score (58) below 0.2 (to exclude likely multiplets). Filtered counts were normalized by scaling each cell to a total of 10,000 counts and log1p-transforming. The 3,000 most variable genes were selected to focus the analysis on biologically informative signals and reduce technical noise. Expression values were scaled to zero mean and unit variance per gene, and dimensionality was reduced by principal component analysis, retaining the first 20 components. A k-nearest-neighbor graph (k = 20) was then constructed in this reduced space and embedded into two dimensions with UMAP for visualization and Leiden clustering (59,60).

Batch integration with Harmony (61) was performed and evaluated, but the uncorrected embedding was retained for the final visualizations. Because the experiment compares biologically distinct conditions (adjacent healthy vs. tumor), integration risks collapsing genuine condition-specific structure together with technical batch effects; the uncorrected representation preserves these between-condition differences more faithfully and was therefore preferred for displaying the contrast between samples.

### *SPP1*, *C1QC*, and *THBS1* signature scores

Within the macrophage compartment, three published transcriptional signatures were used to score functional polarization states at single-cell resolution: the *C1QC*+ and *SPP1*+ tumor-associated macrophage programs described in (17), and the *THBS1*+ macrophage program (47). The *C1QC* signature comprised *C1QA*, *C1QB*, *ITM2B*, *C1QC*, *HLA-DMB*, *MS4A6A*, *CTSC*, *TBXAS1*, *TMEM176B*, *SYNGR2*, *ARHGDIB*, *TMEM176A*, *UCP2*, *CAPZB*, *MAF*, *TREM2*, and *MSR1*; the *SPP1* signature comprised *SPP1*, *PCSK5*, *SLC11A1*, *VCAN*, *SLC25A37*, *FLNA*, *UPP1*, *BCL6*, *AQP9*, *TIMP1*, *VEGFA*, *ADM*, *MARCO*, *FN1*, and *IL1RN*; and the *THBS1* signature comprised *THBS1*, *VCAN*, *S100A12*, *SERPINB2*, *SAP30*, *MEGF9*, *TREM1*, *VEGFA*, *OLR1*, and *PHLDA1*. Per-cell scores were computed with *scanpy.tl.score_genes()* on log-normalized expression. Scores were then compared between conditions (adjacent normal vs. tumor triple co-cultures) using a Student’s t-test on the per-cell score distributions. No multiple-testing correction was applied because only three pre-specified signatures were tested; the resulting p-values were several orders of magnitude below conventional significance thresholds and remain so under Bonferroni adjustment for the three comparisons.

### LIANA+ analysis

Ligand-receptor interactions were inferred with LIANA+ (v1.6) (19) using a differential-expression-based approach, run separately for each condition. Within each condition, differential expression between tumor and adjacent normal samples was computed per cell type using *scanpy.tl.rank_genes_groups()* (default Welch’s t-test), with the opposite condition serving as the reference. The resulting per-cell-type DE statistics (scores, p-values, and adjusted p-values) were passed to *liana. multi.df_to_lr()* together with the condition-specific expression matrix, using the consensus ligand-receptor resource and an expression proportion threshold of 0.1 (interactions retained only if the ligand and receptor were each detected in at least 10% of cells in the relevant cell type). Interactions were ranked by the absolute magnitude of the resulting interaction score, yielding two ranked tables of putative ligand-receptor communications enriched in the tumor and adjacent healthy contexts, respectively. Interactions were filtered for positive interaction scores and adjusted interaction *p*-value < 0.05.

### Publicly available CRC scRNA-seq dataset

To assess the generalizability of the macrophage signature findings, an independent public dataset of colorectal cancer single-cell transcriptomes, Pelka et al.(25), GEO accession (GSE178341, accessed 31 August 2025), was reanalyzed. Cells annotated as macrophages in the original study were selected for analysis. In addition, fibroblast clusters were identified based on expression of the canonical fibroblast markers *LUM*, *DCN*, and *PDGFRA*; clusters that showed positive expression of these markers were retained as fibroblasts. The sample condition (tumor vs. adjacent normal) was taken from the accompanying metadata. Quality filtering required a minimum of 1,500 total counts and a mitochondrial read fraction below 20% per cell; batches with fewer than 20 cells after filtering were removed to ensure stable per-sample estimates. Counts were normalized to 10,000 reads per cell and log transformed. The same *C1QC*, *SPP1*, and *THBS1* macrophage signatures applied to the in-house data were scored on a per-cell basis with *scanpy.tl.score_genes*. To avoid pseudoreplication and respect the biological unit of replication, per-cell scores were aggregated to sample-level means by averaging within each batchID (corresponding to individual patient samples), and each sample was annotated as tumor or adjacent normal. Pairwise relationships between signatures were assessed across samples by Pearson correlation, and differences between tumor and adjacent normal samples were tested with two-sided Student’s t-tests on the sample-level means. Scatter plots of signature pairs colored by condition (with linear fits and Pearson r) and accompanying box-and-strip plots of per-sample scores by condition were generated for the three signature pairs (*C1QC* vs. *SPP1*, *C1QC* vs. *THBS1*, *SPP1* vs. *THBS1*). For visualization, the 3,000 most variable genes were selected, with batchID supplied as the batch key, so that variability was assessed within each patient sample and ranked by consistency across samples rather than dominated by a single donor. Expression was scaled to zero mean and unit variance per gene (clipped at ±10 to limit the influence of extreme values), and dimensionality was reduced by PCA to 20 components. Because this dataset comprises many patient samples processed across distinct batches, batch effects were corrected at the embedding level with Harmony (61). A k-nearest-neighbor graph (k = 20) was constructed on the Harmony-corrected components, and a UMAP embedding (59) was computed from this graph for visualization.

### RNA isolation and quantitative real-time PCR (qPCR)

Total RNA was isolated using the QIAwave® RNA Kit (Qiagen), and cDNA was synthesized with the iScript cDNA Synthesis Kit (Bio-Rad) according to the manufacturer’s protocol. qPCR was performed with Luna Universal qPCR Master Mix (NEB) on a real-time PCR detection system (Bio-Rad). Relative gene expression levels were normalized to human hypoxanthine phosphoribosyltransferase 1 (*HPRT1*). Primer sequences are listed in Table S5. Depending on the dataset and experimental design, statistical significance was assessed using either paired Student’s *t*-tests or one-way ANOVA followed by Dunnett’s multiple comparisons test.

### Bulk RNA sequencing and bioinformatics analysis

Total RNA was isolated as described above. Lysates were additionally treated with DNase I (Qiagen) according to the manufacturer’s instructions. External service providers performed library construction, quality control, and sequencing. Messenger RNA was purified from total RNA using poly-T oligo-attached magnetic beads. Following fragmentation, first-strand cDNA was synthesized using random hexamer primers, followed by second-strand cDNA synthesis using either dUTP for directional libraries or dTTP for non-directional libraries. Libraries were completed by end repair, A-tailing, adapter ligation, size selection, amplification, and purification. Library quality and quantity were assessed using Qubit (Thermo Fisher Scientific), real-time PCR, and the Bioanalyzer (Agilent). Libraries were pooled and sequenced on an Illumina platform based on effective library concentration and the required data output. The clustering of the index-coded samples was performed according to the manufacturer’s instructions. After cluster generation, the library preparations were sequenced on an Illumina platform, yielding paired end reads. Bulk RNA-Seq quantification was performed with kallisto (62) using an index built from the GENCODE/Ensembl transcriptome annotation corresponding to the human reference genome assembly GRCh38 (hg38). Reads were pseudo-aligned to the transcriptome index, and estimated counts and transcripts per million (TPM) values were obtained for each transcript. To enable gene-level differential expression analysis, transcript-level estimates were aggregated to the gene level by summing estimated counts and TPM values across all transcripts annotated to the same gene (simple sum strategy), using the transcript-to-gene mapping derived from the same GENCODE/Ensembl annotation.

### Differential expression analysis and pathway enrichment analysis

Differential expression analysis was performed using the DESeq2 package (63) on raw count data. Genes with low counts were excluded before analysis. Data were subsequently variance-stabilizing transformed (VST) for visualization and exploratory analyses. At the same time, the DESeq2 statistical framework was employed to compute pairwise comparisons between the indicated groups and identify differentially expressed genes. Gene set enrichment analysis (GSEA) was performed using the clusterProfiler package (64) on pre-ranked gene lists derived from DESeq2 differential expression results to assess enrichment for Gene Ontology Biological Processes (GO: BP) and KEGG pathways. Redundancy among enriched GO terms was reduced, when necessary, using semantic similarity-based simplification implemented through the GOSemSim package (65). Mitochondrial pathway enrichment analysis was performed using the fgsea package (66) on pre-ranked gene lists derived from DESeq2 differential expression results. Mitochondrial gene sets were obtained from the MitoCarta3.0 database (67). For visualization, the top enriched pathways were selected based on adjusted *p*-values and displayed according to the normalized enrichment score (NES).

### Isolation of small extracellular vesicles from cell culture media

Small extracellular vesicles (sEVs) were purified from cell culture supernatant (SN) after culturing fibroblasts for 3 days. The HPLM media before cell culture were depleted of extracellular vesicles by ultracentrifugation (Hitachi) at 100,000 × g for 16 hours at 4 °C, followed by ultrafiltration through a 0.22-nm filter. The cell supernatant was collected by pelleting the cells at 300 x g for 5 min, followed by centrifugation at 4000 x g for 20 min. To remove large EVs, 10 ml of the SN was diluted in 20 ml of cold PBS and ultracentrifuged at 30,000 x g for 2 hours at 4 °C, and then the SN was collected as a source of sEVs and pelleted at 100.000 x g for 2 hours, 4 °C. Next, the obtained pellet was washed in cold PBS (100,000 g; 2 hours; 4 °C) and the sEV pellet was subjected to for MACSPlex analysis.

### MACSPlex analysis of small extracellular vesicles (sEVs)

Surface protein profiling of sEVs was performed using the human MACSPlex Exosome Kit (Miltenyi, cat. no. 130-136-851). Briefly, 200 µL sEVs were incubated overnight at room temperature with 7.5 µL MACSPlex Exosome Capture Beads (39 antibody-coated bead types) on an orbital shaker (450 rpm). Samples were then washed with 500 µL MACSPlex buffer and centrifuged at 3000 × g for 5 min to pellet beads. The supernatant was removed, and the bead pellet was subsequently incubated with the MACSPlex Exosome Detection Reagent under continuous shaking. Following detection, beads were washed twice with MACSPlex buffer. Fluorescence signals (FITC, PE, and APC) were acquired using a BD FACSCanto II (BD Biosciences). Data analysis was conducted using FlowJo v10 software. Bead populations were distinguished based on their fluorescence characteristics in the PE and FITC channels, and median APC fluorescence intensity was used as a measure of sEV binding. For each sample, the mean MFI of CD9, CD63, and CD81 was calculated and used as a normalization factor. The MFI of each of the 39 marker-specific bead populations was then divided by this mean value.

### COAD proteome

CPTAC Pan-Cancer COAD proteome data were accessed through the LinkedOmics data portal (https://www.linkedomics.org/data_download/CPTAC-pancan-COAD/) (31) and accessed 23 March 2025. MS2-based protein abundance values were used to visualize TSP1 protein expression between adjacent normal and tumor tissues. Statistical analyses used to assess differences between groups included the Wilcoxon rank-sum test.

### COAD transcriptome and ESTIMATE analysis

The Cancer Genome Atlas Colon Adenocarcinoma (TCGA-COAD) dataset (30) was retrieved from the Genomic Data Commons (GDC) Data Portal (https://portal.gdc.cancer.gov/projects/TCGA-COAD) using the TCGAbiolinks package (68) on 08 August 2025. Tumor samples were subsequently selected, and raw count data were TPM-normalized and log2-transformed before downstream analyses. Further, samples were stratified into *THBS1*-high and *THBS1*-low groups based on the median *THBS1* expression, which was used as input for the ESTIMATE algorithm (29). The Wilcoxon rank-sum test was employed for statistical testing.

### Spatial transcriptomics data visualization and analysis

Publicly available spatial transcriptomics data from colorectal cancer tissue (69) were obtained from the 10x Genomics Visium platform (Visium HD, sample P2 CRC; https://www.10xgenomics.com/platforms/visium/product-family/dataset-human-crc) and accessed 01 May 2026. The data were explored and visualized using the 10x Genomics Loupe Browser (version 9.1.0). Cell clusters were manually annotated using marker genes from precomputed differential expression tables provided in the Loupe Browser, as well as those reported in the original publication. All visualization figures were generated and exported directly from the Loupe Browser.

### BODIPY staining

Macrophages were cultured in µ-Slide 8-well chambers (ibidi) under the indicated conditions for fluorescence staining or harvested and stained in suspension for flow cytometry-based quantification. Quantification of staining was performed in Fiji, as described above. BODIPY™ 493/503 (Thermo Fisher Scientific) was used at a final concentration of 2 nM according to the manufacturer’s instructions, and nuclei were counterstained with DAPI. Flow cytometry was performed on a BD FACSCanto II (BD Biosciences) and analyzed using FlowJo v10 software. One-way ANOVA followed by Dunnett’s multiple comparisons test was used to highlight differences between treated groups and controls.

### FACS analysis

After harvesting cells with Accutase and mechanical scraping, macrophage pellets were washed and resuspended in Zombie/TruStain staining mix (Table S5), then incubated for 10-15 minutes at room temperature (protected from light). Samples were washed, incubated with the antibody cocktail (Table S5) for 20 minutes at room temperature, washed again, and then acquired on a BD FACSCanto II flow cytometer. Data were analyzed using FlowJo v10 software.

### Phagocytosis assay

Phagocytotic activity was measured with pHrodo™ Red *S. aureus* BioParticles™ Conjugate (Thermo Fisher Scientific, cat. no. A10010). Briefly, macrophages were incubated with particles dissolved in complete culture medium at a final concentration of 0.005 mg/mL for 1 hour at 37 °C. Following incubations, macrophages were detached from plates, and the signal was quantified by flow cytometry.

### Seahorse assay

For the measurement of the Oxygen Consumption Rate (OCR) and the Extracellular Acidification Rate (ECAR), the Seahorse XF Mito Stress Test Kit (Agilent, Cat. No. 103015 - 100) was used according to the manufacturer’s recommendation. Cells were seeded in 96-well plates (XFe96/XF Pro Cell Culture Microplate, Agilent). On the day of the Assay, the media was removed. The plate was washed once with Seahorse XF DMEM assay medium (pH 7.4, Agilent, USA) without any supplements. Then 180 µl of Seahorse XF DMEM medium containing 10 mM glucose, 2 mM L-glutamine, and 1 mM pyruvate was added to the wells. The cell plate was incubated for one hour in a CO_2_-free incubator at 37°C. Contents of the Seahorse Mito Stress Test Kit were reconstituted with the Seahorse XF DMEM media to stock concentrations indicated in the Mito Stress Test Kit manual. Sequentially, reagents were added as 10x concentrated solutions to the ports of the Seahorse XFe96/XF Pro sensor cartridges to achieve final concentrations of 1.5 µM oligomycin, 2 µM FCCP, 0.5 µM Rotenone/Antimycin A, and 4 µM Hoechst 33258 (1mg/ml). Following the Seahorse measurement, Hoechst fluorescence was detected in the DAPI channel using the Cytation 5 Cell Imaging Multimode Reader (BioTek, part of Agilent). Collected data was analyzed with the Seahorse Wave Pro Software (10.3.0515, Agilent). OCR levels are displayed per 1.000 cells. Statistical analysis: one-way ANOVA followed by Dunnett’s multiple comparisons test.

### MitoTracker staining and mitochondrial network analysis (MiNA)

For MitoTracker Green (MTG, Cell Signaling Technology, cat. no. FM #9074) staining, macrophages were cultured and treated according to the regimen stated in the figure in µ-Slide 8-well chambers (ibidi). At the indicated time points, cells were washed with 1× PBS and stained with MTG at a final concentration of 200 nM for 30 minutes. Nuclei were counterstained with Hoechst (Invitrogen). Imaging was acquired in Z-stack mode in technical triplicates. Analysis of the mitochondrial network was performed using the MiNA tool (35). Briefly, Z-stacks were converted into orthogonal projections using ZEN 2 EBlue edition software (Zeiss) and subsequently analyzed with the MiNA plugin in Fiji. MiNA-derived parameters were extracted per image/cell. Length- and area-based outputs were converted to µm and µm², respectively, using the original image pixel size. For each donor and condition, MiNA parameters were summarized as mean values and used for downstream statistical analysis. Statistical significance between treated and control macrophage groups was assessed using one-way ANOVA followed by Dunnett’s multiple comparisons test.

### Lipid extraction and UHPLC-MS analysis

Lipids of human primary macrophages were extracted according to Matyash et al (70), with modifications. For extraction, cells were disrupted in 1 ml of methyl tert-butyl ether (MTBE)/methanol (3.33:1, v/v) containing 1 µM butylated hydroxytoluene (BHT). After adding 400 µl ddH2O, phase separation was obtained by centrifugation (13.000 rpm, 5 min, room temperature). The upper organic phase was collected, dried under a N_2_ stream, and reconstituted in 2-propanol/ddH_2_O (1:1, v/v) containing 1% formic acid, 10 mM ammonium formate, and 7.7 µM phosphoric acid for untargeted UHPLC-MS analysis (Infinity II UHPLC; QToF 6560B; Agilent). Lipid samples were analyzed and annotated using MS-DIAL and Skyline. Lipidomics multivariate analysis was done as described in (71). Briefly, normalized lipid intensity values were log2-transformed, autoscaled, and subjected to PLS-DA analysis, followed by calculation of variable importance in projection (VIP) scores.

### Data analysis and visualization

All data processing and analyses were performed in R (version 4.4.2 (2024-10-31)) using RStudio (version 2026.01.1+403). Data visualization was done either in R using ggplot2 or in GraphPad (version 10.5.0 (673)).

## Data availability

The datasets supporting the conclusions of this article are available in the Zenodo repository, https://doi.org/10.5281/zenodo.20273870. Interactive visualization of the single-cell datasets via the Cellxgene browser is available upon request to the corresponding authors.

## Abbreviations

2D: two-dimensional
3D: three-dimensional
ALI: air-liquid interface
Angio-TAMs: proangiogenic tumor-associated macrophages
CAFs: cancer-associated fibroblasts
CD36: cluster of differentiation 36
CD68: cluster of differentiation 68
CL: cardiolipin
CMOs: cholesterol-modified oligos
CRC: colorectal cancer
CSF1R: colony-stimulating factor 1 receptor
CTCF: Corrected Total Cell Fluorescence
HPLM: human plasma-like medium
IFN-TAMs: interferon-primed tumor-associated macrophages
Inflam-TAMs: inflammatory cytokine-enriched tumor-associated macrophages
IntDen: integrated density
LA-TAMs: lipid-associated tumor-associated macrophages
MTG: MitoTracker Green
NES: normalized enrichment score
NFs: normal fibroblasts
PBMCs: peripheral blood mononuclear cells
PC: phosphatidylcholine
PDOs: patient-derived organoids
PDTs: patient-derived tumoroids
PE: phosphatidylethanolamine
PI: phosphatidylinositol
PLS-DA: Partial Least Squares Discriminant Analysis
Prolif-TAMs: proliferating tumor-associated macrophages
PS: phosphatidylserine
Reg-TAMs: immune regulatory tumor-associated macrophages
SSO: Sulfosuccinimidyl oleate
TAMs: tumor-associated macrophages
TG: triacylglycerol
TME: tumor microenvironment
TSP1: thrombospondin 1
VST: variance-stabilizing transformed

## Acknowledgments

The authors would like to thank Sabrina Ladstätter and Christoph Bock for their assistance with single-nuclei library preparation and sequencing. We are grateful to Emine Atas and Lukas Kenner for providing protocols for mitochondrial staining, and to Helga Schachner and Astrid Haase for their support with IHC staining. This research was funded by the Austrian Science Fund (FWF) (10.55776/F8300). I.A. and S.I. were supported by the ERC Synergy grant (KILL-OR-DIFFERENTIATE, 10.3030/856529), Swedish Research Council, Paradifference Foundation, Bertil Hallsten Research Foundation, Cancer Foundation in Sweden, and Knut and Alice Wallenberg Foundation. For open access purposes, the author has applied a CC BY public copyright license to any author-accepted manuscript version arising from this submission. Graphical abstracts and figures in this manuscript were partially created using BioRender.com.

## Author contribution

G.E and K.D. conceptualized the study, administered the project, and drafted the original manuscript. G.E., K.D., M.S., T.W., and S.I. contributed to data interpretation. K.D., I.P., D.V., J.P., M.S., M.H., L.G., A.H., M.M., and P.S. performed the experimental analyses and analyzed the data. K.D. and S.I. conducted computational analyses and data visualization. V.L., G.W., and M.B. provided resources. G.E., I.A., M.S., W.B., and T.W. supervised the study. G.E., T.W., and M.S. acquired funding. All authors reviewed and edited the manuscript.

## Ethics declarations

Healthy donor blood collection in this study was approved by the Ethics Committee of the Medical University of Vienna (approval no. 1513/2025). The use of primary material for the establishment of PDOs/PDTs and NFs/CAFs was approved by the Ethics Committee of the Medical University of Vienna (approval no. 1248/2015). Informed consent was obtained from all participants.

---

**Supplementary Figure 1.**
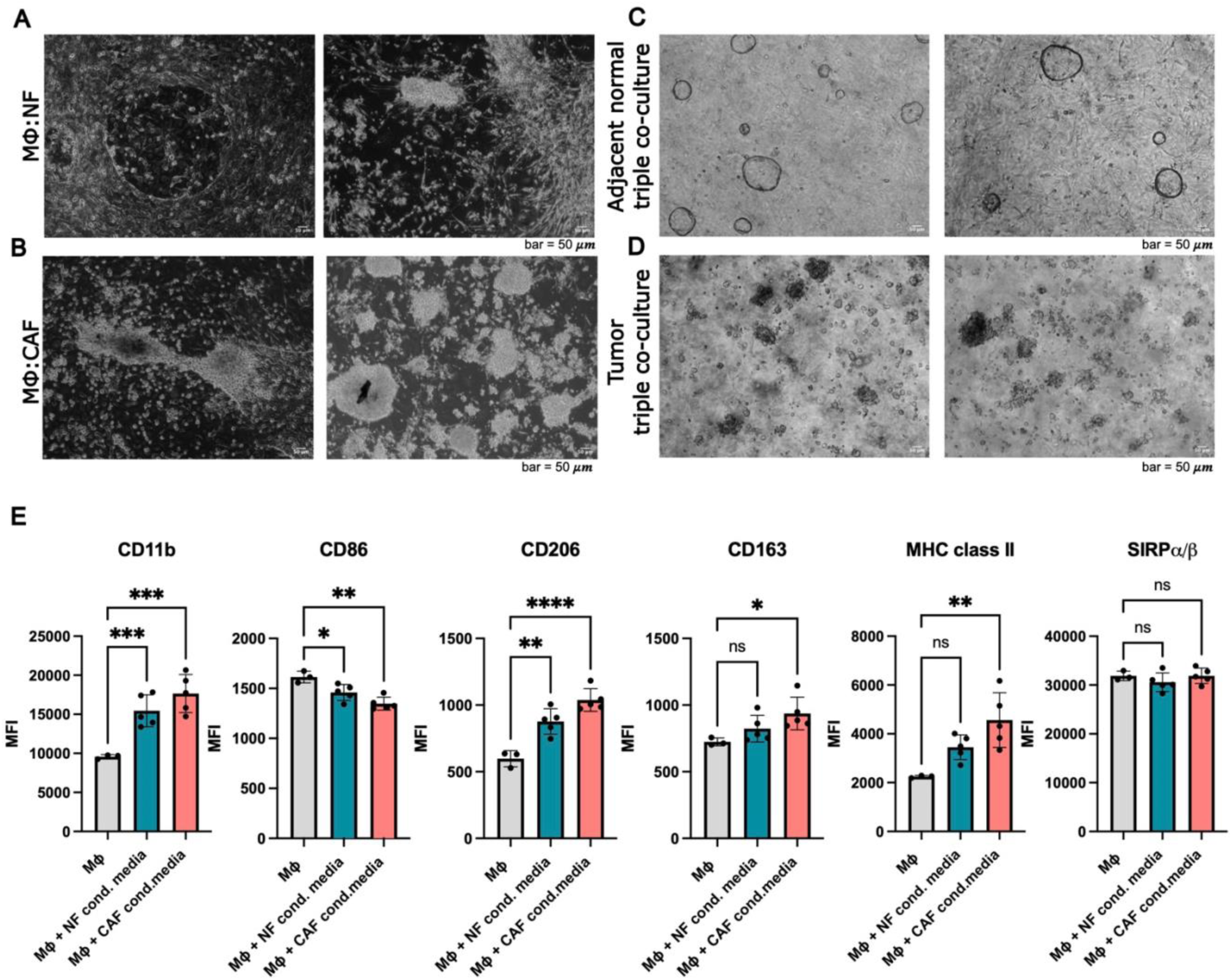
Morphological and immunophenotypic characterization of co-culture models. (A, B, C, D) Brightfield microscopic images of two biological replicates for direct (A) MΦ: NF and (B) MΦ: CAF co-cultures, and (C) adjacent normal and (D) tumor triple co-cultures. (E) Flow cytometry analysis of mean fluorescence intensity (MFI) values for CD11b, CD86, CD206, CD163, MHC class II, and SIRPα/β, comparing marker expression in control macrophages (MΦ) and MΦ cultured in NF-conditioned and CAF-conditioned media (n = 6 tissue-matched CAF and NF pairs). Statistics: one-way ANOVA followed by Dunnett’s multiple comparisons test between control and conditioned media samples (ns, not significant; \**p* < 0.05; \*\**p* < 0.01; *** *p* <0.001; **** *p* < 0.0001).

**Supplementary Figure 2.**
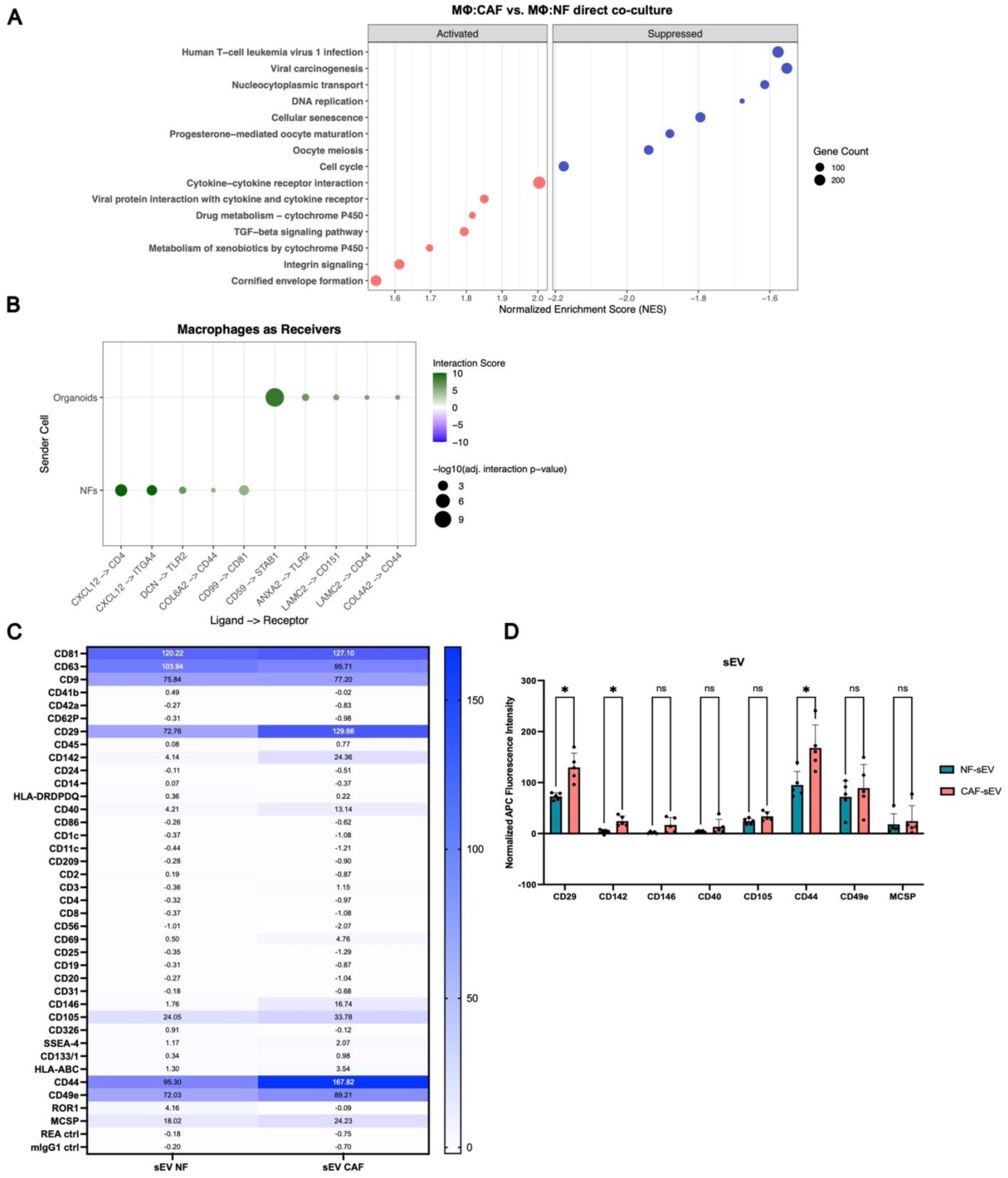
Intercellular communication profiling of macrophage subsets in co-culture models and characterization of fibroblast small extracellular vesicles. (A) GSEA pathway enrichment analysis (GO: KEGG) of bulk RNA-seq data of macrophages cultured in transwell MΦ: CAF vs. MΦ: NF co-cultures. Corresponds to analysis in Fig. 2E. (B) LIANA+ (19) analysis of receptor-ligand interactions inferred from single-nuclei RNA seq data of adjacent normal triple co-cultures, highlighting key predicted signals received by macrophages from either PDOs or NFs. Corresponds to tumor triple co-culture graph in Fig. 2G. (C) Heatmap displaying mean normalized MFI values for CD9, CD63, and CD81, along with 39 individual surface markers of sEVs analyzed by MACSplex between paired NF and CAF samples. (D) The bar chart of selected markers displays the mean ± SD from five matched CAF-NF pairs. Statistics: multiple Student’s t-tests (ns, not significant; \**p* < 0.05).

**Supplementary Figure 3.**
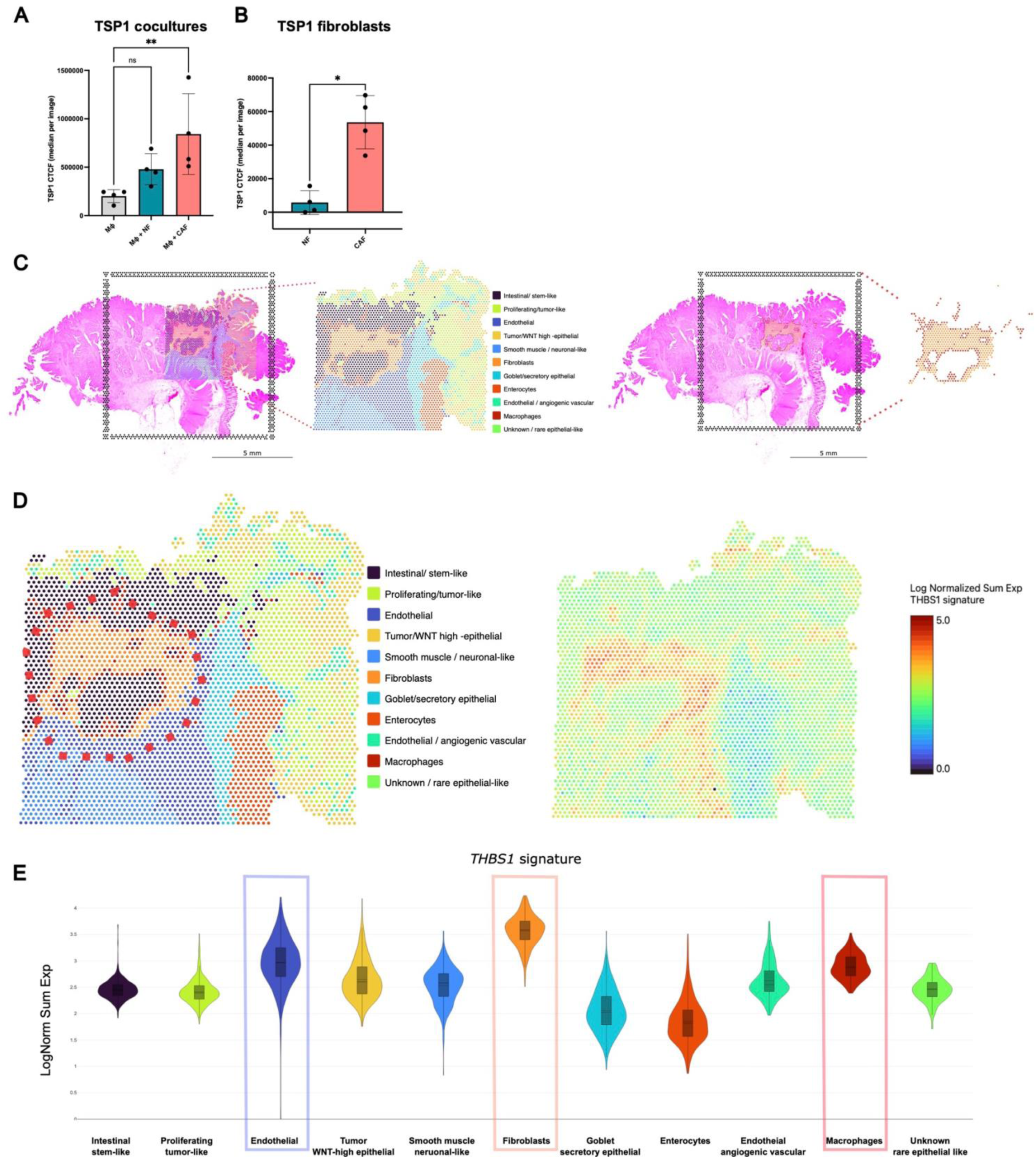
Spatial characterization of *THBS1*expression. (A) Quantification of TSP1 expression in MΦ, MΦ: NF, and MΦ: CAF co-cultures. Fiji/ImageJ- calculated background-corrected total cell fluorescence (CTCF) is presented as the mean ± SD from 4 independent donors, with 3 representative images analyzed per condition. Statistics: one-way ANOVA followed by Dunnett’s multiple comparisons test versus the control group (ns, not significant; \*\**p* < 0.01). (B) CTCF of TSP1 expression in fibroblasts is shown as mean ± SD from four fibroblast pairs, with statistical analysis performed using Student’s t-test (\**p* < 0.05). (C) Spatial transcriptomics analysis of publicly available data (27), displaying a tissue-wide spatial map with manually annotated cell populations (left) and an enlarged region illustrating the structural localization of fibroblast and macrophage populations within the selected tissue area. Data visualization was performed using the 10x Genomics Loupe Browser (version 9.1.0). (D) Zoomed-in view of spatial transcriptomic cell annotations as in (C), red rectangles on the left highlighting the region where the macrophage and fibroblast populations are localized, and spatial feature plot showing log-normalized, summed expression of the *THBS1* gene signature on the right. (E) Violin plot displaying expression of the *THBS1* signature in the indicated cell types annotated from spatial data in (C). Data visualization was performed using the 10x Genomics Loupe Browser (version 9.1.0).

**Supplementary Figure 4.**
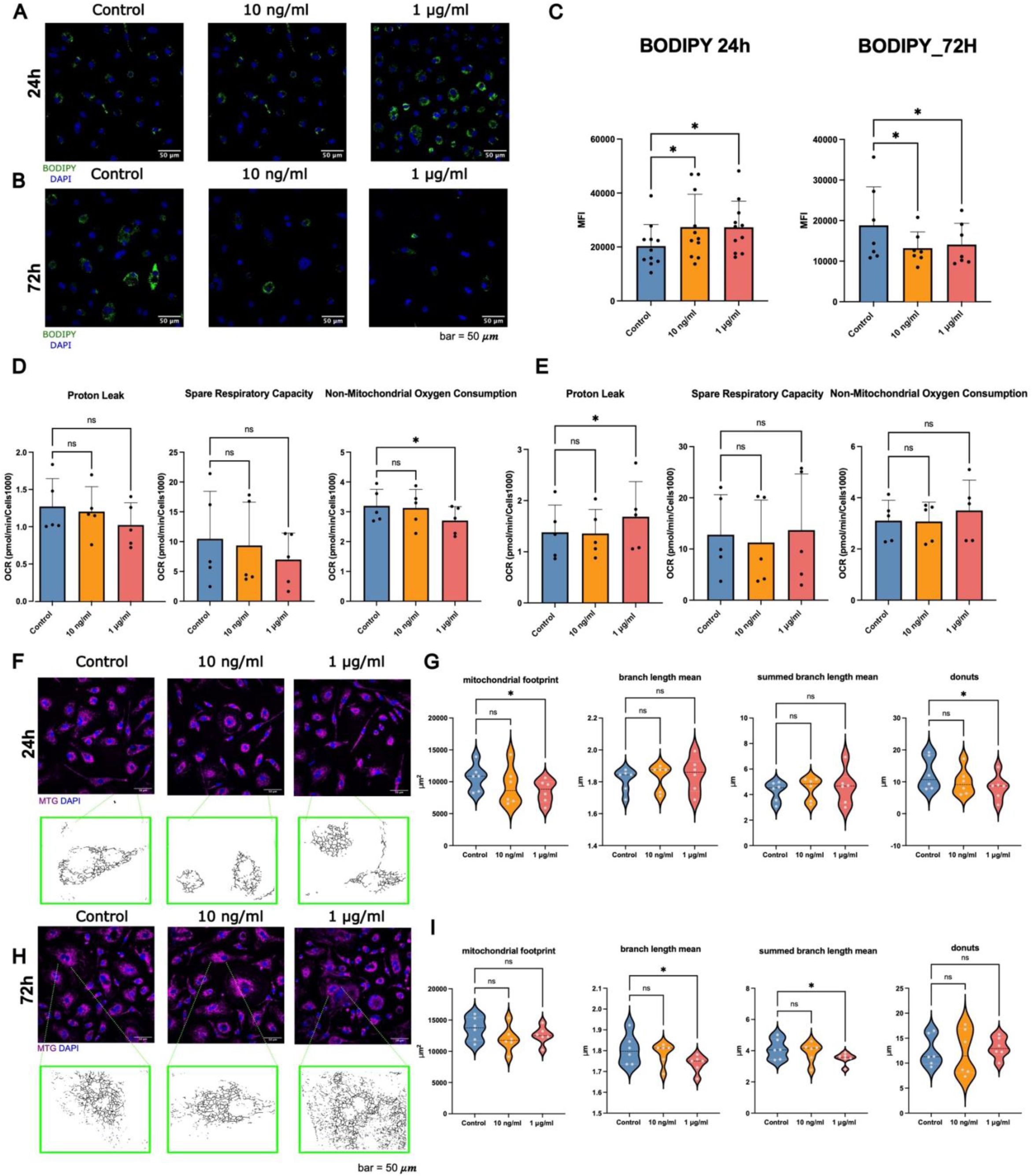
Extended metabolic and mitochondrial analyses during early and late macrophage responses. (A, B) An example of a female donor for BODIPY staining is shown in the graph for (A) 24 hours and (B) 72 hours. (C) BODIPY MFI values quantified by FACS analysis after 24 (left, mean ± SD, n=11) and 72 (right, mean ± SD, n=7) hours of treatment. Statistics: one-way ANOVA followed by Dunnett’s multiple comparisons test versus the control group (\**p* < 0.05). (D, E) Additional parameters from Seahorse® MitoStress tests for (D) 24 or (E) 72 hours of rTSP1 treatment of macrophages correspond to the main Fig. 5F and H. Statistics: one-way ANOVA followed by Dunnett’s multiple comparisons test versus the control group (ns, not significant; \**p* < 0.05). (F, H) An example of a female biological replicate for MTG staining is shown in the graph for 24 and 72 hours. (G, I) Results of parameters derived from MiNa analysis for (G) 24 and (I) 72 hours of treatment (mean ± SD, n=6), corresponding to the main Fig. 5K and L. Statistics: one-way ANOVA followed by Dunnett’s multiple comparisons test versus the control group (\**p* < 0.05; \*\*\**p* < 0.001; ns, not significant).

**Supplementary Figure 5.**
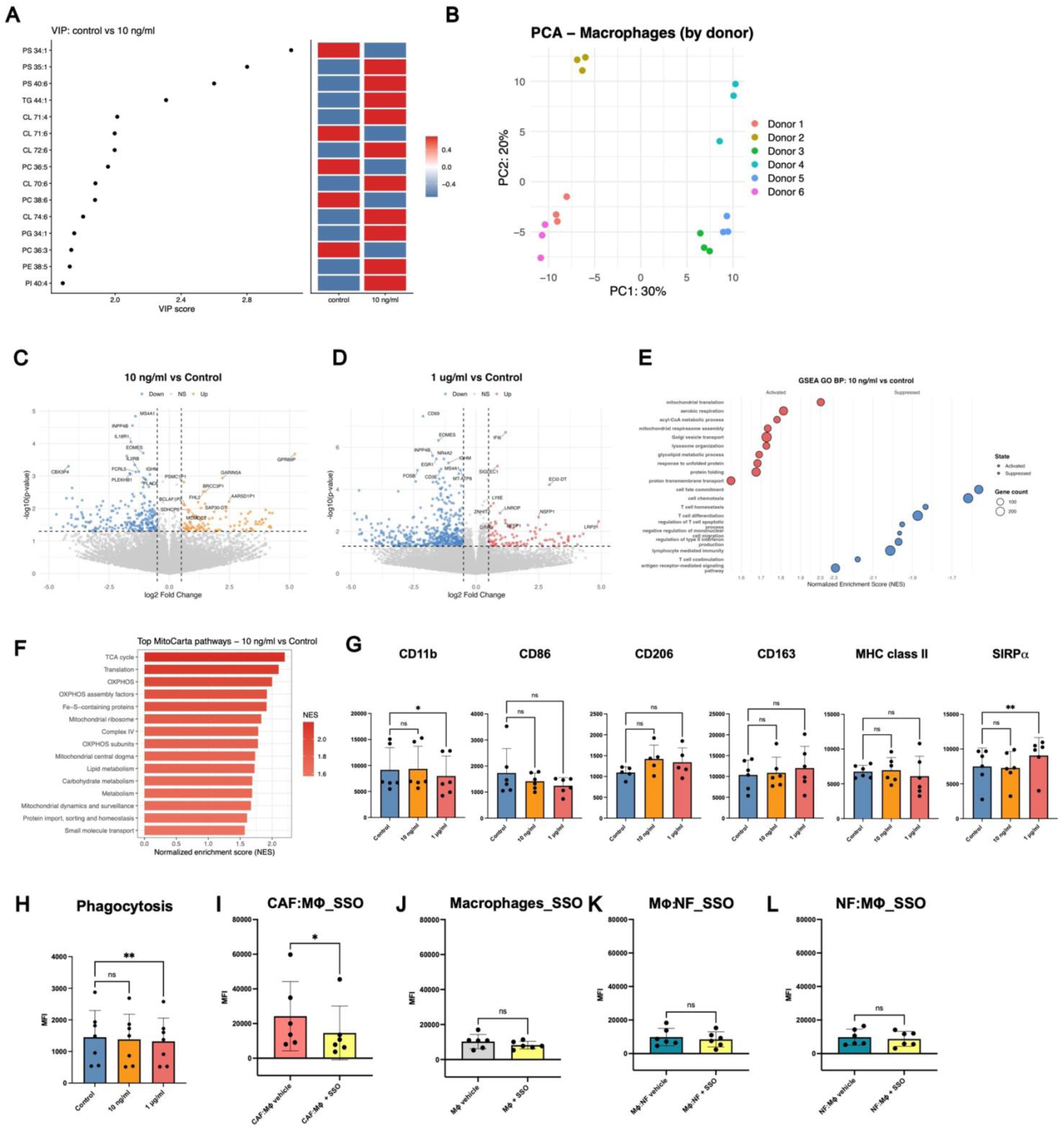
Additional lipidomic, transcriptomic, and functional analyses of stimulated macrophages. (A) Heatmap showing enrichment of VIP scores of different lipid classes inferred from lipidomic analyses in 10 ng/ml rTSP1-treated versus control macrophages. PS, phosphatidylserine; TG, triacylglycerol; CL, cardiolipin; PC, phosphatidylcholine; PE, phosphatidylethanolamine. (B) PCA plot demonstrating donor-dependent clustering of macrophages used from bulk RNA-seq, corresponds to main Fig. 6C. (C, D) Volcano plots from DESeq2 analysis of bulk RNA-seq data of macrophages treated with (C) 10 ng/ml and (D) 1 μg/ml rTSP1 versus control. Differentially expressed genes are highlighted: upregulated (red and orange) and downregulated (blue), defined by |log2fc| ≥ 0.5 and *p*-value < 0.05. (E) GSEA of GO: BP pathways in macrophages treated with recombinant TSP1 (10 µg/ml) compared to control. (F) MitoCarta 3.0 pathway enrichment analysis of rTSP1 (10 ng/ml)-treated macrophages versus control, with bar color indicating both magnitude and direction of enrichment. (G) Flow cytometry immunophenotype analysis showing MFI values for CD11B, CD86, CD206, CD163, MHC Class II, and SIRPα/β in control, 10 ng/ml, and 1 μg/ml-treated macrophages (mean ± SD, n=5). Statistics: one-way ANOVA followed by Dunnett’s multiple comparisons test versus the control group (not significant; \**p* < 0.05, \*\**p* < 0.01). (H) Flow cytometric quantification of phagocytic uptake using pHrodo™ Red *S. aureus* BioParticles™ in untreated control versus rTSP1-treated macrophages (mean ± SD, n=7). Statistics: one-way ANOVA (ns, not significant; \*\**p* < 0.01). (I, J, K, L) Flow cytometry quantification of BODIPY staining: (I) CAFs co-cultured with MΦ and treated with vehicle (DMSO) or SSO (50 μM); (J) MΦ monocultured and treated with vehicle (DMSO) or SSO (50 μM); (K) MΦ co-cultured with NFs and treated with vehicle (DMSO) or SSO (50 μM); and (L) NFs co-cultured with MΦ and treated with vehicle or SSO (50 μM) (mean ± SD, n=6). Statistics: Student’s t-test (\**p* < 0.05).

